# Ca^2+^/Calmodulin Dependent Protein Kinase Kinase-2 (CaMKK2) promotes Protein Kinase G (PKG)-dependent actin cytoskeletal assembly to increase tumor metastasis

**DOI:** 10.1101/2023.04.17.536051

**Authors:** Debarati Mukherjee, Rebecca A Previs, Corinne N Haines, Muthana Al Abo, Patrick K Juras, Kyle C Strickland, Binita Chakraborty, Sandeep Artham, Regina Whitaker, Katherine L Hebert, Jake Fontenot, Steven R Patierno, Jennifer A Freedman, Frank H Lau, Matthew Burow, Ching-Yi Chang, Donald P. McDonnell

## Abstract

Triple-negative breast cancers (TNBCs) tend to become highly invasive early during cancer development. Despite some successes in the initial treatment of patients diagnosed with early-stage localized TNBC, the rate of metastatic recurrence remains high with poor long-term survival outcomes. Here we show that elevated expression of the serine/threonine-kinase, Calcium/Calmodulin (CaM)-dependent protein kinase kinase-2 (CaMKK2), is highly correlated with tumor invasiveness. We determined that genetic disruption of CaMKK2 expression, or inhibition of its activity, disrupted spontaneous metastatic outgrowth from primary tumors in murine xenograft models of TNBC. High-grade serous ovarian cancer (HGSOC), a high-risk, poor-prognosis ovarian cancer subtype, shares many genetic features with TNBC, and importantly, CaMKK2 inhibition effectively blocked metastatic progression in a validated xenograft model of this disease. Probing the mechanistic links between CaMKK2 and metastasis we defined the elements of a new signaling pathway that impacts actin cytoskeletal dynamics in a manner which increases cell migration/invasion and metastasis. Notably, CaMKK2 increases the expression of the phosphodiesterase PDE1A which decreases the cGMP-dependent activity of protein kinase G1 (PKG1). This inhibition of PKG1 results in decreased phosphorylation of Vasodilator-Stimulated Phosphoprotein (VASP), which in its hypophosphorylated state binds to and regulates F-actin assembly to facilitate contraction/cell movement. Together, these data establish a targetable CaMKK2-PDE1A-PKG1-VASP signaling pathway that controls cancer cell motility and metastasis. Further, it credentials CaMKK2 as a therapeutic target that can be exploited in the discovery of agents for use in the neoadjuvant/adjuvant setting to restrict tumor invasiveness in patients diagnosed with early-stage TNBC or localized HGSOC.

## Introduction

Breast cancer recently surpassed lung cancer as the leading cause of cancer incidence, with an estimated 2.3 million new cases in 2020, representing approximately 11.7% of all cancer cases worldwide^1^. It remains the most common cause of cancer-associated deaths among women, and is responsible for nearly 15% of all new cancer cases each year in the United States^2^. Approximately, 90% of all breast cancer related deaths can be attributed to metastasis^3^. While the 5-year overall survival rate of breast cancer patients without metastasis is now over 80%^4^, this figure drops to ∼ 25% for patients diagnosed with metastatic disease^5^. Furthermore, 12-17% of newly diagnosed cases are classified as triple-negative breast cancer (TNBC), reflecting the absence of expression of hormone or growth factor receptors^6^. TNBC tumors tend to become highly invasive early during cancer development and are biologically more aggressive with worse prognosis than other breast tumor subtypes. Women with early stage TNBC who present with locally invasive, lymph node-positive disease are at a high risk of developing distant metastatic recurrence within 2-3 years from initial diagnosis. Despite advances in treatments for patients with early stage, locally advanced TNBC, most do not achieve a pathologic complete response (pCR) and 5-year recurrence rates remain ∼80% ^7 8 9^. The high rate of distant recurrence, even among women presenting with early-stage disease, reinforces the urgent need to understand the fundamental molecular mechanisms driving localized tumor cell invasion and distant metastasis in TNBC. It is anticipated that elucidation of the mechanisms underlying these processes will allow the identification of interventions that can significantly impact the invasiveness and metastatic potential of TNBC.

Calcium/Calmodulin (CaM)-dependent protein kinase kinase 2 (CaMKK2) is a serine/threonine-kinase that couples a variety of external stimuli to intracellular pathways involved in the regulation of cell proliferation, survival and metabolism^10^. This enzyme was initially identified as a component of the Calcium/Calmodulin-dependent protein kinase (CaMK) cascade that is activated in response to elevated intracellular Ca^2+^. However, several studies have also demonstrated that CaMKK2 can exhibit significant activity independent of Ca^2+^/Calmodulin (CaM) binding (autonomous activity)^10 11^. The most well characterized downstream effectors of CaMKK2 activity include CaMKI, CaMKIV and AMPKα^10 12^. CaMKK2 acts through CaMKI to activate ERK and CREB which in turn directly impacts neurite outgrowth in neuroblastoma and osteoclast differentiation during bone development^13–15^. Further, CaMKK2-CaMKI can regulate cytoskeletal architecture and actin-based motility in developing neurons^16^. CaMKK2-dependent activation of CaMKIV directly controls liver cancer cell proliferation^17^, and modulates the homeostatic functions of brain-derived serotonin^18^. AMPKα functions as a key integrator/regulator of CaMKK2 activity, and this signaling axis regulates metabolic responses to energetic stress, particularly in the brain^19^, liver^20^ and adipose tissue^21^. In addition to its role in maintaining whole body energy homeostasis, the CaMKK2-AMPK pathway has been shown to integrate cellular responses to cellular Ca^2+^ influxes and in this manner modulate anoikis^22^, autophagy^23 24^, cell viability/survival^25^, micropinocytosis^26^ and cellular migration^27^.

CaMKK2 is aberrantly overexpressed in a number of cancers, and its role in primary tumor models of prostate cancer^28^, lung cancer^22^, glioblastoma^29^, ovarian cancer^30^, renal cell carcinoma^24^ and pancreatic cancer^26^ have been explored. Further, we recently reported that CaMKK2 is strongly expressed within cancer cells and in the stroma of all types of human breast tumors and that its overexpression is directly correlated with more aggressive TNBC^31^. Although CaMKK2 within myeloid cells was shown to support tumor growth by reprogramming the tumor immune microenvironment, its cancer cell intrinsic roles and how they impact cancer metastasis have not been elucidated. Interestingly, CaMKK2 was recently implicated as an upstream regulator of actin polymerization and stress fiber assembly in osteoclasts^32^. The presence of a robust network of thick intracellular actin stress fibers is a hallmark of migrating cells and provides the force necessary to trigger cellular motility^33^. Indeed, CaMKK2 action has previously been shown to regulate actin-based cellular motility in developing neurons^16^. These observations, together with the finding that CaMKK2 is aberrantly overexpressed in highly metastatic tumors^31^, prompted us to probe if and how CaMKK2 action regulates tumor cell motility, migration and metastatic dissemination from primary tumors. We find that depletion of CaMKK2 (or its pharmacological inhibition) disrupted actin cytoskeleton organization in cancer cells which resulted in decreased spontaneous metastasis in murine xenograft models of cancer. This work highlights the potential utility of recently developed, highly specific, CaMKK2 inhibitors as treatments for TNBC or HGSOC or to prevent metastatic recurrence of early-stage cancers.

## Results

### Elevated expression of CaMKK2 is associated with aggressive phenotypes and poor outcomes in patients with basal subtypes of breast cancer

Previously, we determined that the expression of CaMKK2 was elevated in invasive TNBC^31^ and now report that higher expression of this enzyme is associated with reduced overall survival in patients with the basal subtype of breast tumors (**Figure 1A; *top left***). No differences in overall survival among patients with either luminal or HER2+ disease was observed. Basal tumors are immunohistochemically defined by the lack of expression of the estrogen receptor (ER), progesterone receptor (PR) or HER2 with TNBCs representing a large fraction of cancers exhibiting the basal-like molecular subtype^34^. Interestingly, validated gene signatures predicting heightened tumor cell motility and invasiveness^35^ and spontaneous metastatic potential from solid tumors^36^ were found to be associated with elevated CaMKK2 expression in tumors from patients with basal-like or luminal disease (**Figures 1B, C**). Furthermore, a well-validated 6-gene signature that predicts the occurrence of lung metastasis from primary breast tumors^37^, was found to be significantly upregulated in basal tumors with high CaMKK2 expression (**Figure 1D**). Finally, CaMKK2-high basal tumors (but no other tumor subtypes) showed significantly heightened aggregate expression of a well-validated molecular signature predicting micrometastatic outgrowth from the primary tumor in breast cancer (**Figure 1E**)^38^. Taken together, these results suggest that increased expression of CaMKK2 in primary tumors may be a key determinant of increased invasiveness and metastatic capacity in breast cancer, especially in patients with hormone receptor-negative, basal subtype(s) of this disease.

**Figure 1.**
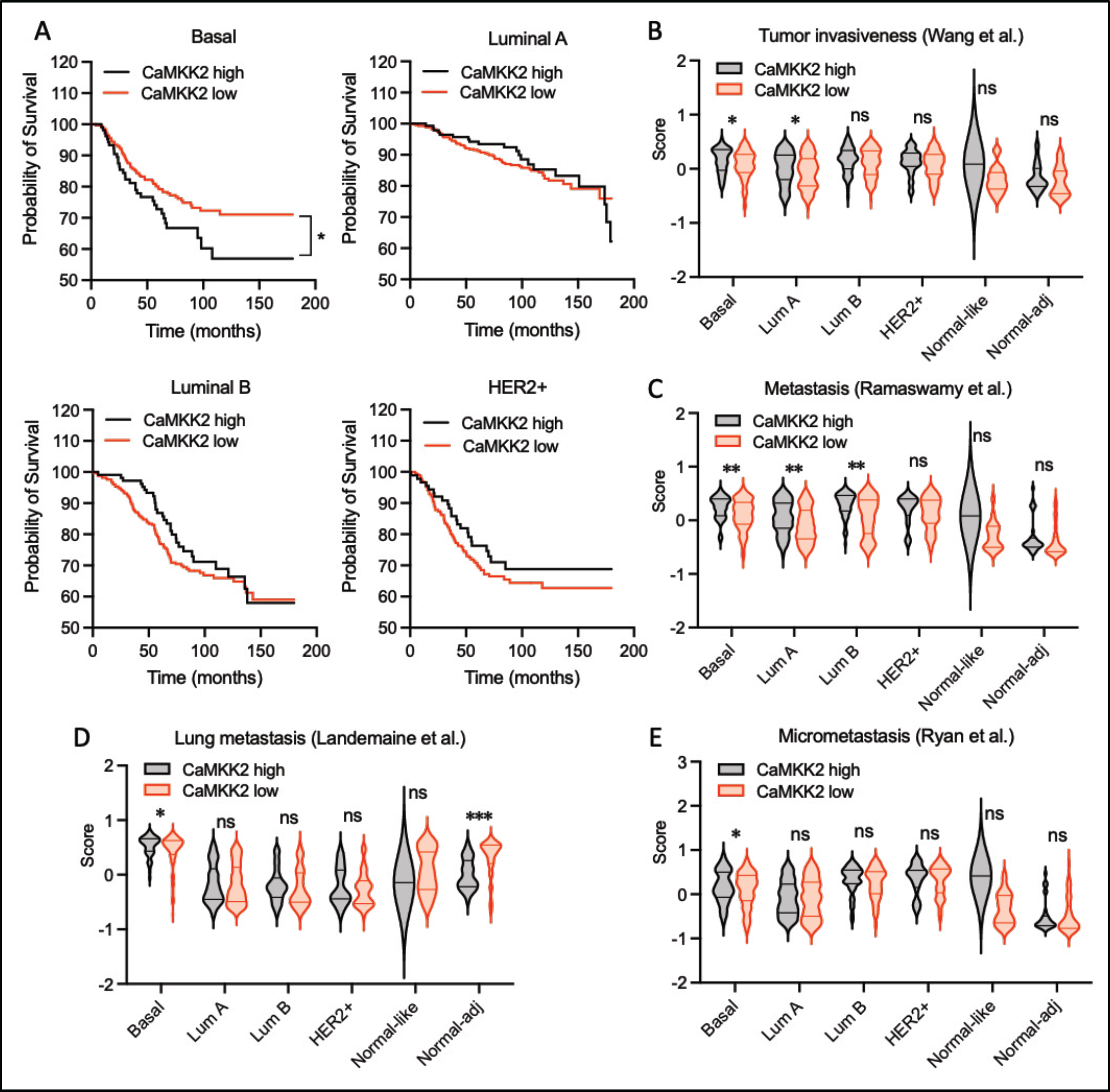
Clinical relevance of CaMKK2 in metastatic TNBC. **A,** Kaplan-Meier plots showing the association between *CaMKK2* level and the overall survival of patients with basal (n=431), luminal A (n=362), luminal B (n=439) or Her2+ (n=596) tumor subtypes. Patients were grouped into CaMMK2-high or –low by upper quartile (lower or higher than 75^th^ percentile). The x-axis shows time in months, and the y-axis shows the overall survival probability. Plots were generated using KM-plotter^100^ (https://kmplot.com), **B-E,** Violin plots depicting the score for the gene signatures published in the indicated articles^35–38^. Samples were grouped into CaMMK2-high or –low by quartiles (equal or less than 25^th^ percentile, and equal or greater than 75th percentile). The x-axis shows the different breast cancer subtypes, and the y-axis shows the score between -1 and 1. Data are expressed as the mean ± SEM. *P < 0.05, **P < 0.01, ***P < 0.005, and ****P < 0.001, by Student’s t test **(B, C, D** and **E**) and log-rank test **(A**)

### Depletion of CaMKK2 impairs the migratory potential of invasive breast cancer cells

To directly probe the cancer cell intrinsic role(s) of CaMKK2 expression on tumor cell invasiveness and metastasis in TNBC, we assessed the impact of inhibiting CaMKK2 expression and activity in three cellular models of this disease: MDA-MB-231, BT-20 and HCC1954. Changes in AMPK phosphorylation was used as a biomarker to assess the degree of CaMKK2 inhibition achieved. Transient shRNA-dependent knockdown of CaMKK2 expression in all three cell lines (**Figures 2A, B, Supplementary Figure 1A**) significantly inhibited cell migration when compared to controls (**Figures 2C, D; Supplementary Figure 1B**). Similar outcomes were observed in cells treated with STO-609 (a commercially available CaMKK2 inhibitor), or GSK1901320 (GSKi) (a highly selective CaMKK2 inhibitor)^31^ (**Figures 2E, F; Supplementary Figure 1C**). Furthermore, inhibition of CaMKK2 activity negatively impacted the invasiveness of both MDA-MB-231 (**Figure 2G**) and BT-20 cells (**Figure 2H**). Additionally, loss of CaMKK2 expression or activity also inhibited the migration of HOC7 cells, a cellular model of ovarian cancer, indicating that the role of CaMKK2 in supporting cancer cell motility and migration is not restricted to breast cancer **(Supplementary Figures 1D, E**).

**Figure 2.**
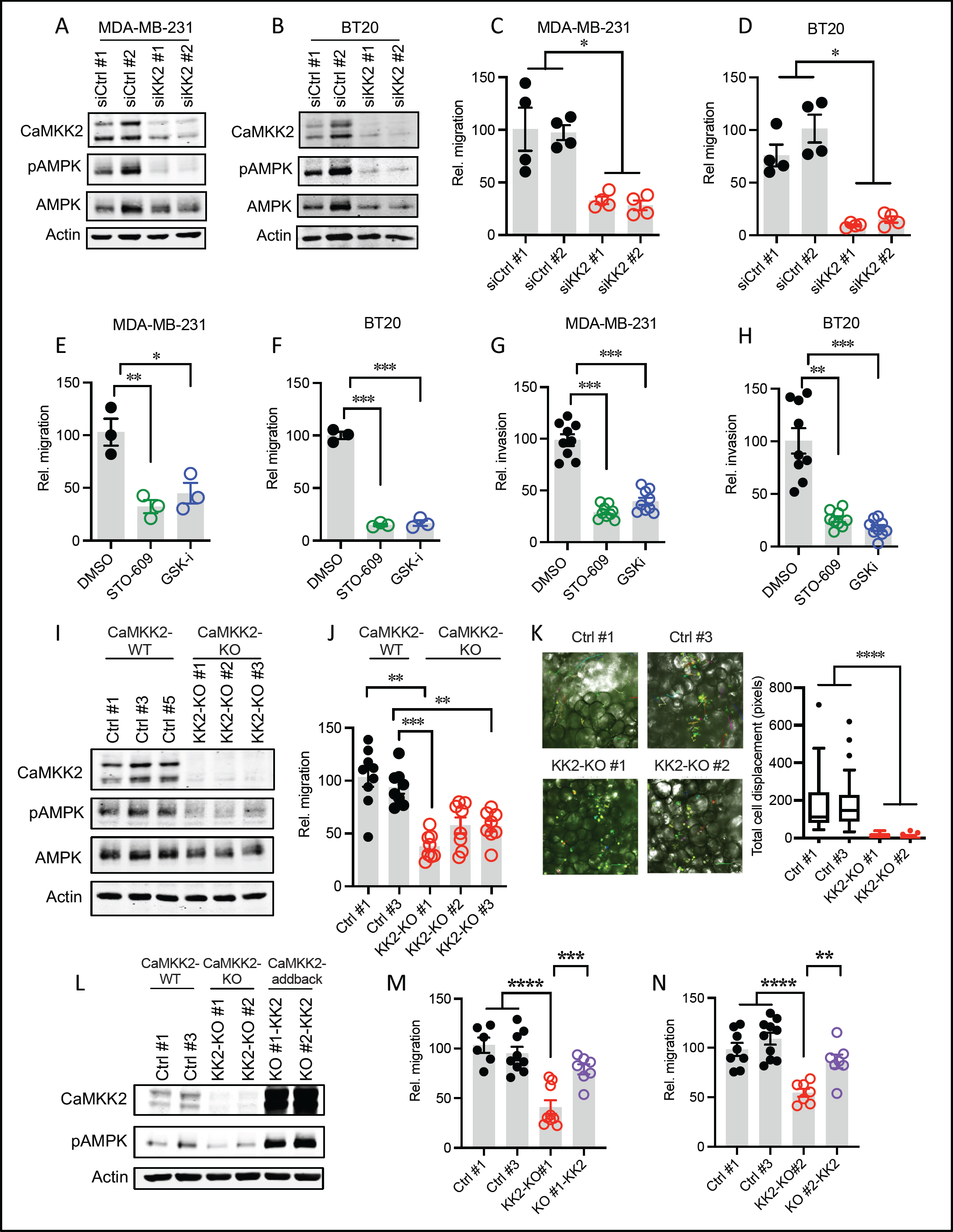
Ablation of CaMKK2 impairs migration in invasive breast cancer cells. **A,** Representative blots showing knockdown of CaMKK2 in highly invasive MDA-MB-231 and **B,** BT-20 TNBC cells. MDA-MB-231 and BT-20 cells were transfected with either control siRNAs (siCtrl #1, siCtrl #2) or siRNAs targeting CaMKK2 (siKK2 #1, siKK2 #2). 48 hours post transfection, cell lysates were harvested and western immunoblots were used to analyze CaMKK2 and phospho-AMPK⍺ (T172) expression levels. β-Actin was used as a loading control. Representative resultsfrom two independent experiments are shown. **C,** CaMKK2 knockdown impaired migratory ability in MDA-MB-231 cells and, **D,** BT-20 cells *in vitro.* Migration assays were performed with MDA-MB-231 and BT-20 cells transfected with either control siRNAs (siCtrl #1, siCtrl #2) or CaMKK2 siRNAs (siKK2 #1, siKK2 #2). Data are plotted as mean ± SEM; n = 6 random fields measurements from a total of four individual experiments. **E,** Pharmacological inhibition of CaMKK2 with STO-609 (10µM, 48h) and GSKi (1µM, 48h) resulted in impaired migration in MDA-MB-231 and, **F,** BT20 cells *in vitro.* Data are plotted as mean ± SEM; n = 9 random fields measurements from a total of three transwell chambers. **G,** Pharmacological inhibition of CaMKK2 with STO-609 (10µM, 48h) and GSKi (1µM, 48h) resulted in reduced invasiveness in MDA-MB-231 and, **H**, BT20 cells *in vitro*. Data are plotted as mean ± SEM; n = 4-5 random fields measurements from two individual invasion chambers, **I**, Representative blots confirming CRISPR-mediated knockout of CaMKK2 in MDA-MB-231 cells. (see also **Supplementary Figure 1H**), **J,** CRISPR-mediated genetic ablation of CaMKK2 leads to impaired migratory ability in all three CaMKK2-KO clones of MDA-MB-231 cells *in vitro*, Data are plotted as mean ± SEM; n = 4-5 random fields measurements from two individual transwell chambers. **K,** Depletion of CaMKK2 reduced cellular movement in MDA-MB-231 cells when cultured with fresh human breast tissue. Fluorescence microscopy and bright field images were captured every 5 minutes over 6 hours to track cellular movement (see also **Supplementary Videos 1-4**). **L,** Representative blots showing addback of CaMKK2 expression in CaMKK2-KO clones #1 and #2. Western immunoblots were used to analyze CaMKK2 and phospho-AMPK⍺ expression levels. β-Actin was used as a loading control. **M, N,** Stable overexpression of CaMKK2 rescues migratory capability in both CaMKK2 KO clones #1 and #2. Data are plotted as mean ± SEM; n = 4-5 random fields measurements from two individual transwell chambers. *P < 0.05, **P < 0.01, ***P < 0.005, P values were calculated using unpaired Student’s t test.

To directly assess the role of CaMKK2 in migration and metastatic potential, we used CRISPR/Cas9-based genome editing to stably knock out CaMKK2 expression in highly metastatic MDA-MB-231 cells. Four separate sg-RNAs (sg-RNA #A, #B, #C and #D) targeting specific sequences at the 5’ end of the CaMKK2 coding region were used (**Supplementary Figure 1F, *in red***). Interestingly, these sgRNAs failed to abolish the lower band of the CaMKK2 doublet that is found in most cells (**Supplementary Figure 1G**). We identified a potential cryptic translation initiation sequence ∼150bp downstream from the primary start site. Using sg-RNAs designed to specifically target sequences downstream from this site (sgRNA #1, #2 and #3) (**Supplementary Figure 1F, *in green***), we were able to successfully abolish both CaMKK2 variants (**Supplementary Figure 1H**). Following clonal expansion, a single CaMKK2-KO clone was randomly selected corresponding to each of the three sgRNAs #1, #2 and #3. Loss of CaMKK2 expression and downstream function (AMPK phosphorylation) was confirmed in all three CaMMK2-KO cell clones (hereafter referred to as KK2-KO #1, KK2-KO #2 and KK2-KO #3) (**Figure 2I**). Consistent with what was seen in the CaMKK2-silenced cells, all three CaMKK2-KO clones exhibited significantly reduced migratory capability when compared to control clones (**Figure 2J, Supplementary Figures 1I, J**). This was confirmed in 3D experiments using the Sandwiched White Adipose Tissue (SWAT) preparation technique^39 40^ whereby two control clones and two CaMKK2-KO clones were mixed with freshly minced human breast tissue (adipocytes) to mimic the three-dimensional, multicellular microenvironment in the human breast and the cancer cell movement was observed directly via time-lapse fluorescence microscopy over 6 hours. Loss of CaMKK2 significantly hindered cellular movement in both CaMKK2-KO clones when cultured in fresh human adipocytes (**Figure 2K, Supplementary Videos 1-4**). To determine the extent to which CaMKK2 is necessary to maintain cancer cell migration *in vitro,* we stably overexpressed this protein in two CaMKK2-KO cell lines, KK2-KO #1 and KK2-KO #2 (**Figure 2L**). Overexpression of CaMKK2 rescued the impaired migratory capacity in the knockout clones (**Figures 2M, N**). Taken together, these data indicate that sustained CaMKK2 expression is required to maintain the migratory phenotype of TNBC cells and may ultimately control their metastatic potential.

### CaMKK2-deficiency impairs metastatic outgrowth from primary tumors *in vivo*

To directly assess the impact of CaMKK2 on metastatic outgrowth, three CaMKK2 KO clones of luciferase-labeled MDA-MB-231 cells (KK2-KO #1, KK2-KO #2 and KK2-KO #3) and three non-specific control cell variants (Ctrl #1, Ctrl #3 and Ctrl #5) were orthotopically grafted into the mammary fat pad of immunodeficient mice. The primary tumors were allowed to grow until they reached a maximum size of approximately 2000 mm^3^, following which, the secondary organs were excised and the extent of metastasis in secondary organs was examined by *ex vivo* bioluminescence imaging (**Figure 3A**). We observed that CaMKK2 deficiency significantly affected the metastatic behavior of MDA-MB-231 cells. Indeed, metastasis to secondary organs was found to occur at significantly lower levels compared to that of controls (**Figures 3B, C, D**), resulting in dramatically reduced overall metastatic burden in all three groups bearing CaMKK2-KO tumors (**Figure 3E**). In contrast, no consistent effect on primary tumor growth was observed when CaMKK2 was depleted (**Supplementary Figure 2A).** This indicates that, rather than controlling tumor growth, CaMKK2 likely impacts breast tumor pathobiology by controlling metastatic dissemination from the primary tumor.

**Figure 3.**
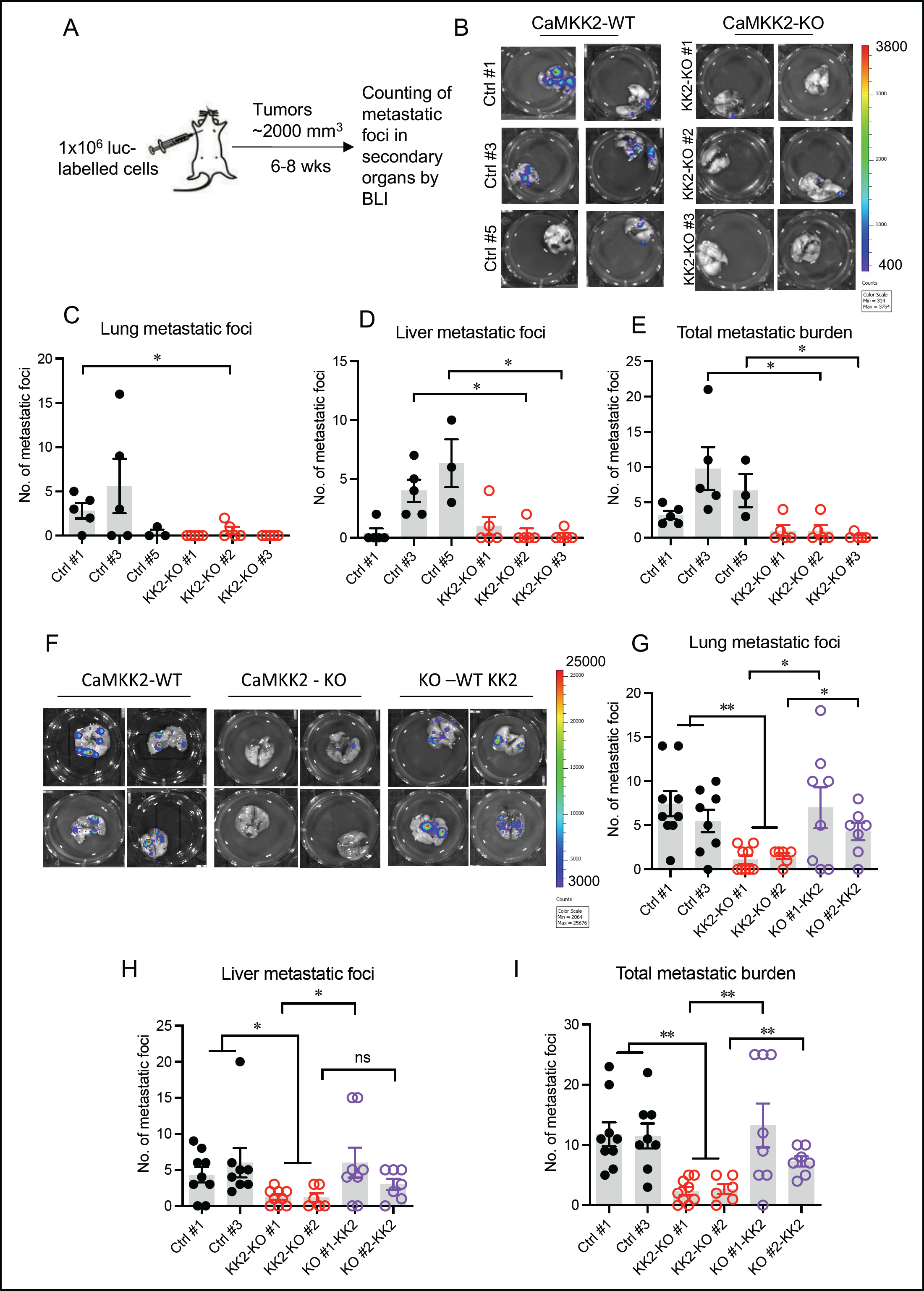
Genetic ablation of CaMKK2 impairs metastasis from the primary tumor *in vivo*. **A,** Schematic representation of experimental protocol. 6-week old nude mice were orthotopically injected in the mammary fat pad with three control clones, Ctrl #1 (n=5), Ctrl #3 (n=5), Ctrl#5 (n=3) and three CaMKK2-KO clones, KK2-KO #1 (n=5), KK2-KO #2 (n=5) and KK2-KO #3 (n=5). The tumors were allowed to reach a volume of 2000 mm^3^. The mice were then sacrificed and the lung and liver metastatic burden was immediately analyzed *ex vivo.* **B,** Representative BLI images showing metastatic foci in the lung in mice bearing a tumor of approx. 2000 mm^3^. **C,** Quantitation of the number of lung metastatic foci in mice injected with either control clones or CaMKK2 knock out clones as observed by *ex vivo* bioluminescence imaging. **D,** Quantitation of the number of metastatic foci in the liver. **E,** Quantitative analyses of the total metastatic burden (metastatic foci in the liver and lung taken together for each individual animal). **F,** 6-week old female nude mice were orthotopically injected in the mammary fat pad with two control cell clones, Ctrl #1 (n=9), Ctrl #3 (n=8); two CaMKK2-KO cell clones, KK2-KO #1 (n=9), KK2-KO #2 (n=6) and the CaMKK2 KO cell clones with re-expression of CaMKK2, KO #1-KK2 (n=8) and KO #2-KK2 (n=7). The tumors were allowed to reach a volume of approx. 2500 mm^3^. The mice were then sacrificed and the lung and liver metastatic burden was analyzed *ex vivo.* Representative BLI images showing metastatic foci in the lung in mice bearing a tumor of approx. 2500 mm^3^ was visualized. **G**, Quantitation of the number of lung metastatic foci. **H,** Quantitation of the number of metastatic foci in the liver. **I**, Quantitative analyses of the total metastatic burden (includes metastatic foci in the liver and lung taken together for each individual animal). Data are expressed as the mean ± SEM. \**P <* 0.05, \*\**P <* 0.01, \*\*\**P <* 0.005, and \*\*\*\**P <* 0.001, by Student’s *t* test.

Clinically detectable metastasis is the endpoint of a series of sequential cell-biological events, starting with the initial migration of tumor cells from the primary site, invasion into the surrounding adipose tissue, resulting ultimately in vascular access^41^. The latter steps of the metastatic cascade include extravasation into secondary organs where the disseminated tumor cells that survive may seed the growth of secondary tumors in a metastasis-receptive niche^5 41^. Having established that intratumoral ablation of CaMKK2 significantly impairs spontaneous metastasis, we next determined at which point CaMKK2 impacts the metastatic cascade. To this end we inoculated the CaMKK2-KO cells directly into the circulation of immunocompromised mice via intravenous injection bypassing the initial steps of the metastatic cascade. Intriguingly, depletion of CaMKK2 in cells directly injected into the circulation had no significant impact on the rate of metastatic outgrowth (**Supplementary Figure 2B**). These data suggest CaMKK2 expression likely impacts overall metastatic progression by positively influencing the initial stages of metastasis, including local migration/invasion of tumor cells from the primary site into the tumor-associated stroma.

To confirm that sustained CaMKK2 expression is necessary for spontaneous metastasis from the primary tumor, we orthotopically grafted CaMKK2-KO cells having stable re-expression of the protein (KO#1-KK2 and KO#2-KK2) into the mammary fat pad of immunocompromised mice. We found that overexpression of CaMKK2 in both knockout clones was accompanied by increased spontaneous metastatic outgrowth in secondary organs and increased total metastatic burden (**Figures 3F, G, H, I**). Interestingly, animals bearing tumors in which CaMKK2 was re-expressed exhibited increased tumor burden when compared to animals bearing CaMKK2-depleted tumors (**Supplementary Figure 2C**). Taken together, our findings indicate that CaMKK2 is a key driver of metastatic progression and controls the metastatic potential of the primary tumor.

Pharmacological intervention with either STO-609 or GSKi had no impact on primary tumor growth (**Figure 4A**), but significantly reduced spontaneous metastasis from the primary tumor (**Figures 4B, C, D**). These findings are consistent with the results of the genetic studies described above and, together, justify further exploration/optimization of CaMKK2 inhibitors to decrease the invasiveness and metastatic potential of mammary tumors.

**Figure 4.**
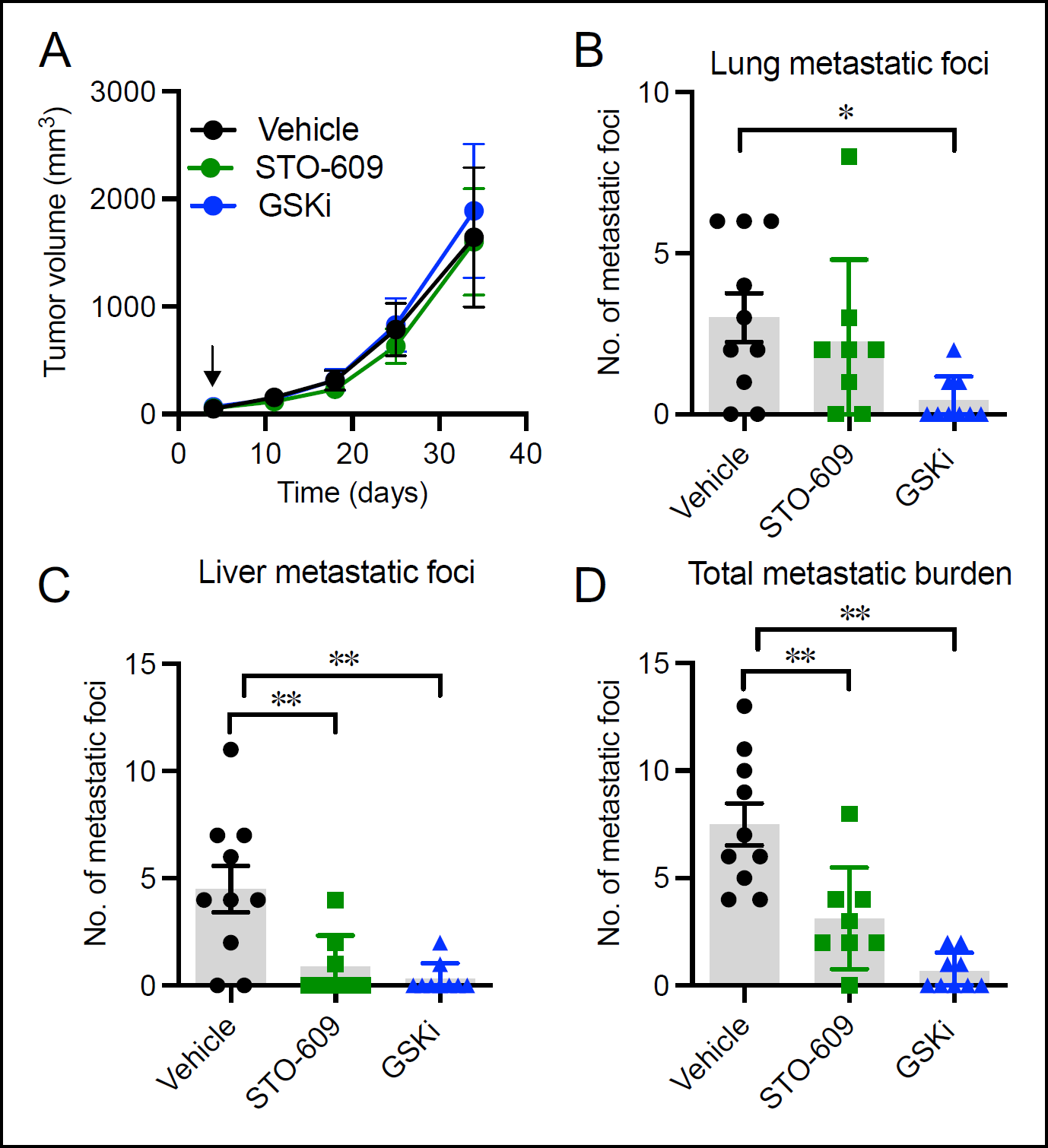
Pharmacological inhibition of CaMKK2 impairs metastasis from the primary tumor *in vivo*. **A,** MDA-MB-231 cells were orthotopically injected into the mammary fat pad of 6-week old female nude mice. The mice were then randomized (five days post tumor cell injection, as indicated) and dosed with Vehicle (n=10), STO-609 (30mg/kg) (n=8) or GSKi (10mg/kg) (n=9) by intraperitoneal injections every 3^rd^ day. Tumor growth rate was monitored by calipers twice weekly. The tumors were allowed to reach a volume of 2000 mm3. **B**, Quantitative analysis of the number of lung metastatic foci in mice dosed with Vehicle, STO-609 or GSKi. The mice were sacrificed after the primary tumor reached a volume of 2000 mm^3^. The lung metastatic foci were analyzed and enumerated *ex vivo*. **C**, Quantitation of the number of metastatic foci in the liver. **D,** Quantitative analyses of the total metastatic foci in the lung and liver taken together. Data are expressed as the mean ± SEM. \**P <* 0.05, \*\**P <* 0.01, \*\*\**P <* 0.005, and \*\*\*\**P <* 0.001, by 2-way ANOVA followed by Bonferroni’s multiple-correction test (**A** or unpaired Student’s *t* test (**B, C** and **D**).

### Ablation of CaMKK2 negatively impacts cytoskeletal assembly and cancer cell motility through enhanced VASP phosphorylation

A major characteristic of migrating cells is the presence of intracellular filamentous actin structures (stress fibers), which generate the traction forces necessary to propel cells forward^33 42 43^. Actin-based stress fibers can be divided into three main groups: “dorsal stress fibers” are thick non-contractile actin filament bundles that elongate radially towards the cell center through vasodilator-stimulated phosphoprotein (VASP)-driven actin polymerization at focal adhesions, “transverse arcs” are thin actin filament bundles that undergo retrograde flow towards the cell center to ultimately fuse to form thick contractile “ventral stress fibers” that drive tail retraction and cell shape changes in migrating cells^44^. While dorsal stress fibers drive directional migration^33^, ventral stress fibers are the major force-generating actin structures in migrating cells^45^. Given the impact of CaMKK2 inhibition on cell migration and motility, we proceeded to examine whether depletion of CaMKK2 in

MDA-MB-231 cells caused changes in actin polymerization and stress fiber assembly within these cells. Visualization of F-actin by fluorescent phalloidin staining revealed an almost complete loss of thick dorsal and ventral stress fibers in the CaMKK2-KO cells compared to control cells (**Figure 5A, Supplementary Figure 3**). This finding is similar to that of a previous study which demonstrated in osteosarcoma cells, that knockdown of CaMKK2 expression or STO-609-mediated inhibition of CaMKK2 activity resulted in defective actin stress fiber assembly^32^. Interestingly, the abnormal stress fiber phenotype in CaMKK2-depleted MDA-MB-231 cells was accompanied by a concomitant increase in the phosphorylation of VASP (**Figure 5B).** VASP promotes F-actin polymerization by localizing to regions of dynamic actin remodeling, to drive actin stress fiber assembly and migration at the leading edge of the cell^46–48^. This protein is a downstream substrate of cAMP-dependent and cGMP-dependent protein kinases (PKA and PKG) which directly phosphorylate VASP at Serine 157 and Serine 239, respectively^49–51^. Serine 239 site is adjacent to the VASP G-actin binding site (residues 234-237)^52^ and its phosphorylation inhibits its ability to bind actin and interferes with actin polymerization^53^. Notably, PKG preferentially phosphorylates VASP at the S239 site, but subsequently exhibits some residual activity at the S157 residue^49 54^. We observed markedly increased levels of VASP phosphorylation at Serine 239 in CaMKK2-depleted cells which was reinforced by treatment of cells with a calcium-specific ionophore, ionomycin (**Figure 5C**). CaMKK2-depletion had a lesser impact on Serine 157 phosphorylation, while the Serine 239 residue was highly phosphorylated in CaMKK2-deficient cells, compared to the controls. Of note, ionomycin-induced Ca^2+^ uptake in control cells did not result in the activation of AMPK (a biomarker for CaMKK2 activity), suggesting that in this context the responses to CaMKK2 activation occurred independently of calcium in MDA-MB-231 cells. Pharmacological inhibition of CaMKK2 activity also increased phosphorylation of VASP at Serine 239 in both MBA-MB-231 (**Figure 5D**) and BT-20 cells (**Figure 5E**). The increase in VASP phosphorylation, in response to CaMKK2 inhibition, tracked with reduced migration potential with or without ionomycin-induced Ca^2+^ influx in MDA-MB-231 cells (**Figure 5F**). Similar observations were made in BT-20 cells (**Figure 5G**). Together, these data suggest that inhibition of CaMKK2 results in increased phosphorylation of VASP to disrupt stress fiber assembly and inhibit TNBC cell motility.

**Figure 5.**
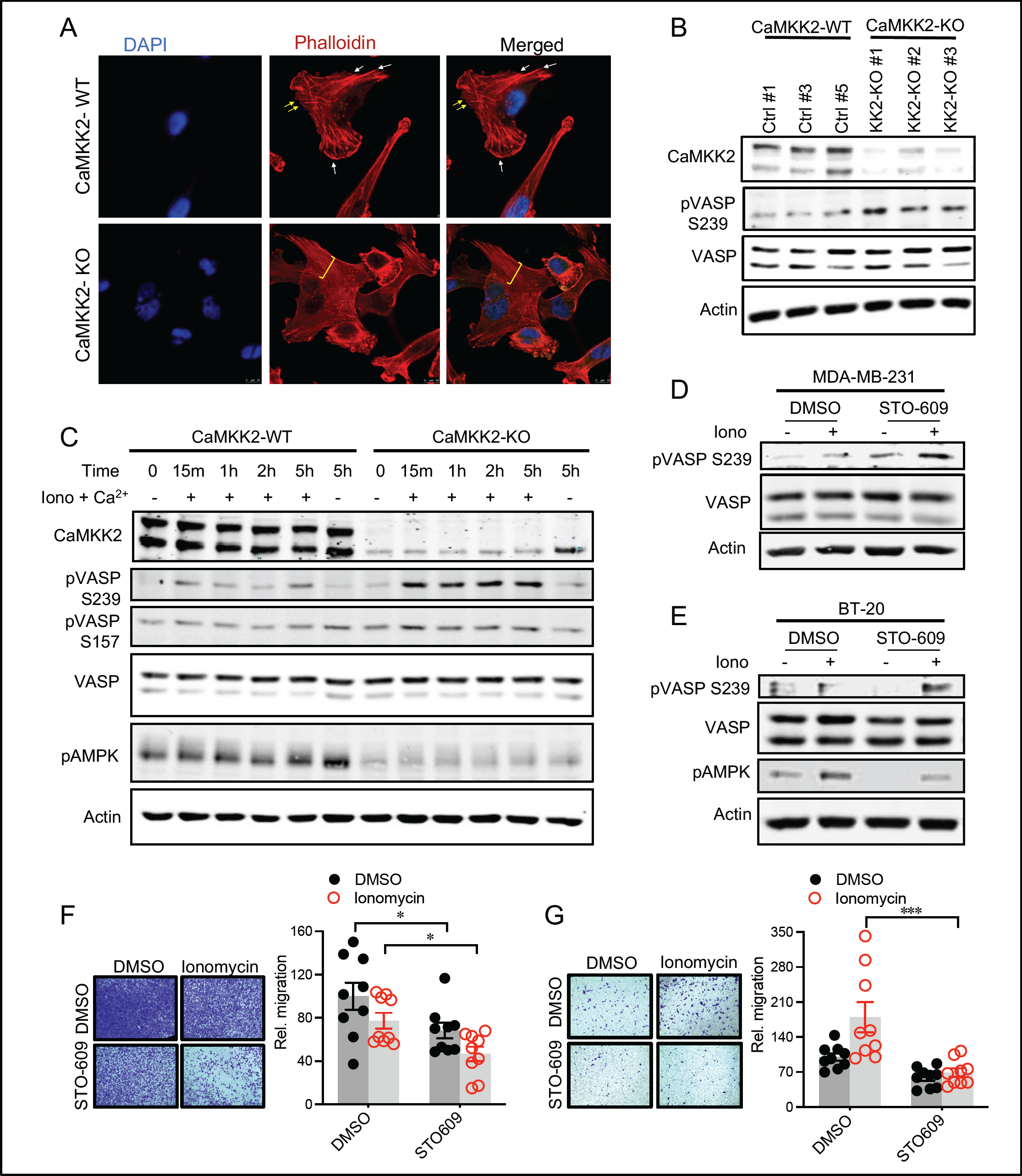
Depletion of CaMKK2 leads to increased phosphorylation of VASP at the Serine 239 residue resulting in impaired cytoskeletal assembly and cell motility. **A,** Representative images showing impaired cytoskeletal assembly in CaMKK2-KO cells as compared to control cells. Visualization of F-actin by phalloidin staining in CaMKK2-WT (Ctrl #1) and CaMKK2-KO (KK2-KO #1) cells revealed almost complete loss of thick ventral stress fibers in cells lacking CaMKK2 expression. Yellow arrows, dorsal stress fibers; white arrows, ventral stress fibers; yellow bracket, transverse arcs. The scale bar represents 10 μm. **B**, Representative blot showing increased phosphorylation of VASP at the Serine 239 residue in all the CaMMK2 KO cell clones as compared to the control clones. Cell lysates were harvested and western immunoblots were used to analyze CaMKK2, phospho-VASP (Ser 239) and total VASP expression levels. β-Actin was used as a loading control. Representative results from two independent experiments are shown. **C**, Representative blot showing VASP phosphorylation at the Serine 239 residue is further enhanced in CaMKK2-KO cells in response to ionomycin (1μM). Cells were then treated with either DMSO or ionomycin (1μM) with CaCL_2_ (1mM) for varying time periods (as indicated) before harvesting for immunoblot analysis. Representative results from three independent experiments are shown. **D-E**, Pharmacological inhibition of CaMKK2 (STO-609, 10μM) increases phosphorylation of VASP (especially with ionomycin (1μM, 2h)) in metastatic MDA-MB-231 cells (D) and BT20 (E). The cells were treated with either DMSO or STO-609 for 48 hours before being exposed to either DMSO or ionomycin (1μM) with CaCL_2_ (1mM) for 2 hours. Representative blots of phospho-VASP vs. total VASP are shown. Representative results from three independent experiments are shown. **F-G**, Inhibition of CaMKK2 results in decreased migratory ability of MDA-MB-231 (**F**) and BT20 (**G**) cells *in vitro.* Cells were treated with either DMSO or STO-609 for 24 hours in serum-rich media followed by 24 hours of treatment in serum-starved media. The cells were then exposed to either DMSO or ionomycin (1μM) with CaCL_2_ (1mM) for 2 hours. Cells were then harvested and migration assays were performed. Data are plotted as mean ± SEM; *n* = 4-5 random fields measurements from two individual experiments. *P < 0.05, **P < 0.01, ***P < 0.005, *P* values were calculated using unpaired Student’s *t* test.

### Inhibition of PKG1 restores the migratory capability of CaMKK2-deficient cells

The molecular mechanism(s) by which CaMKK2 regulates VASP phosphorylation was next assessed. Considering that PKG1 is the only upstream kinase that phosphorylates VASP at Serine 239, we hypothesized that its increased activity in CaMKK2-depleted cells may be responsible for the inhibitory phosphorylation of VASP (**Figure 6A**). To explore this possibility, we assessed the effects of targeting PKG1 activity in CaMKK2-deficient cells using a highly specific, substrate competitive small molecule inhibitor of this enzyme (RKRARKE)^55^. We demonstrated that inhibition of PKG1 activity reduced VASP phosphorylation levels in TNBC cells in which CaMKK2 was inhibited pharmacologically (**Figures 6B, C**) or genetically (**Figures 6D, E**). The ability of RKRARKE to reverse the impact of CaMKK2 inhibition/depletion on VASP phosphorylation implicated PKG1 in the pathway linking CaMKK2 to cell motility. This was confirmed by showing that pre-treatment of cells with RKRARKE restored the migratory phenotype of CaMKK2-KO cells to that of wild-type cells (**Figure 6F**), and rescued the migration noted in STO-609-treated MDA-MB-231 cells (**Figure 6G**). Similar results were found using KT5823, an allosteric inhibitor of PKG1^56–58^ (**Figures 6H, I**). Collectively, our data suggest that decreased expression/activity of CaMKK2 in breast cancer cells results in increased PKG1 activity, and that this increases VASP phosphorylation to inhibit stress fiber assembly, ultimately impairing cellular motility and migratory capability.

**Figure 6.**
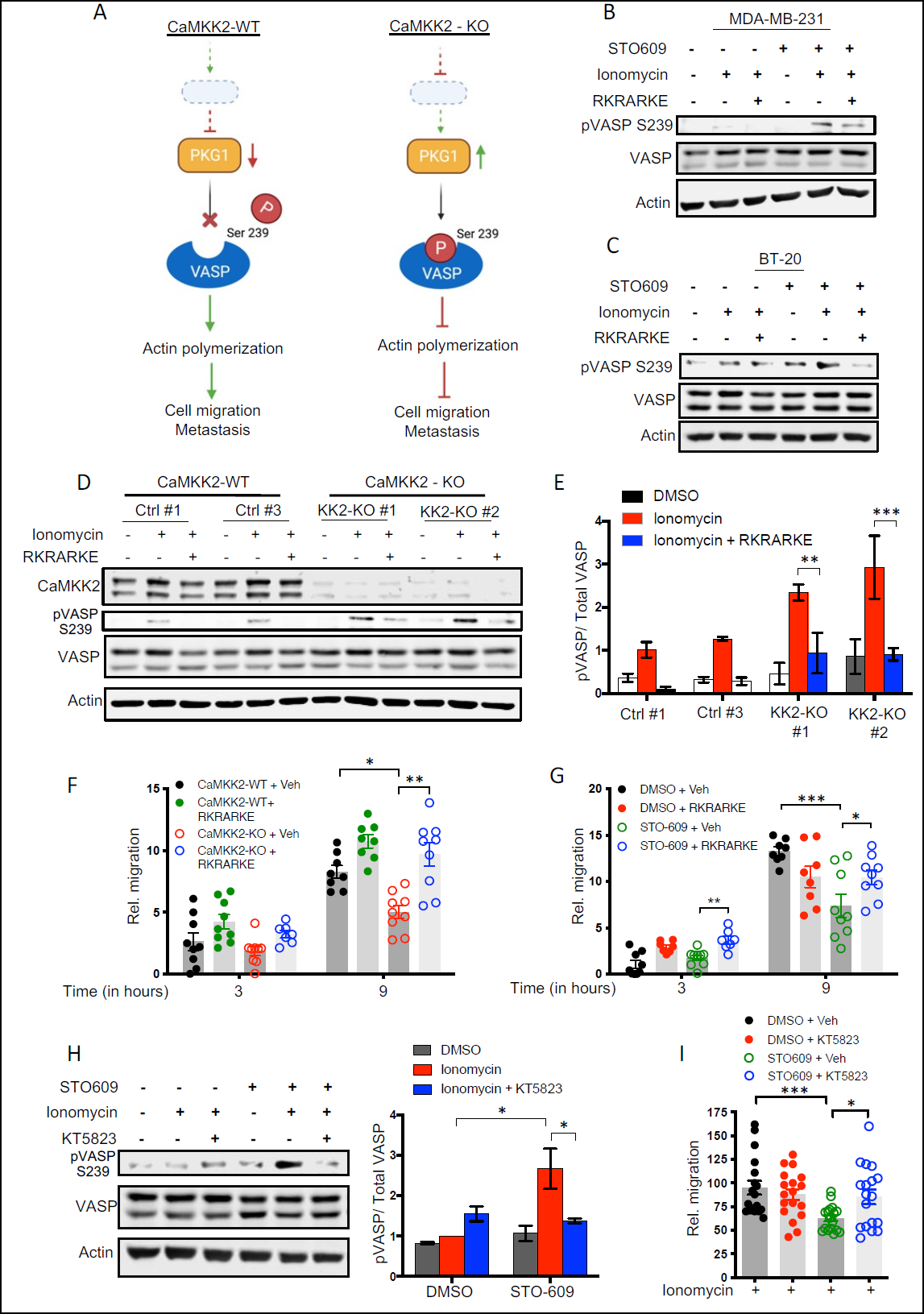
Inhibition of PKG1 blocks phosphorylation of VASP in CaMKK2-depleted cells and restores cellular migration *in vitro*. **A,** Schematic representation of working hypothesis explaining the mechanism by which CaMKK2 may be driving cell migration and metastatic dissemination. **B, C,** Pretreatment with RKRARKE (competitive inhibitor of PKG1) (100µM, 2.5h) in CaMKK2-inhibited cells results in decreased VASP phosphorylation in parental MDA-MB-231 and BT-20 cells, respectively. Cells were treated with either DMSO or STO-609 for 48 hours then with RKRARKE for 30 mins before being exposed to DMSO or ionomycin (1µM) with CaCL_2_ (1mM) for 2 hours (total treatment time with RKRARKE equals 2.5 hours). Representative blots from three independent experiments **(B**) and one experiment **(C**) are shown. **D, E,** Inhibition of PKG1 with RKRARKE blocks phosphorylation of VASP at the Serine 239 residue in CaMKK2-KO cells. CaMKK2-WT (Ctrl #1) and CaMKK2-KO (KK2-KO #1) cells were pretreated with RKRARKE for 30 mins before being exposed to DMSO or ionomycin (1µM) with CaCL_2_ (1mM) for 2 hours. Representative blots from two independent experiments **(D**) and quantitative results for the band intensities of phospho-VASP vs. total VASP are shown **(E**). Data are plotted as mean ± SEM from two independent experiments. **F,** RKRARKE (100µM, 2.5h) pre-treatment in CaMKK2-KO cells restores the migratory phenotype in these cells. CaMKK2-WT and CaMKK2-KO cells were cultured in serum-starved media overnight and then pretreated with RKRARKE for 30 mins before being exposed to ionomycin (1µM) with CaCL_2_ (1mM) for 2 hours. Cells were then harvested and transferred to transwell chambers and allowed to migrate for 3 hours and 9 hours, as indicated. Data are plotted as mean ± SEM; *n* = 4-5 random fields measurements from two individual transwells. **G,** Inhibition of PKG1 with RKRARKE (100µM, 2.5h) pretreatment can rescue the migratory ability of CaMMK2-inhibited cells. Cells were treated with either DMSO or STO-609 for 24 hours in serum-rich media followed by 24 hours of treatment in serum-starved media. The cells were then pretreated with RKRARKE for 30 mins before being exposed to ionomycin (1µM) with CaCL_2_ (1mM) for 2 hours. Cells were then harvested and migration assays were performed for the time periods indicated. Data are plotted as mean ± SEM; *n* = 4-5 random fields measurements from two individual transwells. **H,** Pretreatment with KT5823 (highly selective allosteric inhibitor of PKG1) (5µM, 24h) in CaMKK2-inhibited cells results in decreased VASP phosphorylation in MDA-MB-231 cells. Cells were pretreated with either Vehicle or KT5823 overnight before being exposed to DMSO or ionomycin, as indicated for 2 hours. Representative blots (*left*) and quantitative analysis of band intensities (*right*) from two independent experiments are shown. **I,** KT5823 pre-treatment (5µM, 16h) of CaMKK2-inhibited cells restores the migratory phenotype. MDA-MB-231 cells were treated with either DMSO or STO-609 for 24 hours in serum-rich media followed by 24 hours of treatment in serum-starved media with either Vehicle or KT5823. The cells were then exposed to ionomycin (1µM) with CaCL_2_ (1mM) for 2 hours. Cells were then harvested and allowed to migrate for 9 hours. Data are plotted as mean ± SEM; *n* = 4-5 random fields measurements from four individual transwells. *P < 0.05, **P < 0.01, ***P < 0.005, *P* values were calculated using unpaired Student’s *t* test.

### CaMKK2 negatively regulates the expression of a key upstream modulator of PKG1 activity

It has been established that cGMP is the primary positive regulator of PKG1 activity and thus we posited that CaMKK2 may regulate pathways/processes which control the cellular pool of this second messenger. To this end we evaluated whether the expression or activity of specific 3’,5’-cyclic nucleotide phosphodiesterases (PDEs) which hydrolyze cGMP, modulate PKG1 activity and downstream VASP phosphorylation in metastatic breast cancer cells. The human PDEs comprise a superfamily of 11 enzymes that hydrolyze either cAMP or cGMP. We focused on the PDEs that specifically hydrolyze cGMP (PDE5, PDE6 and PDE9) or which prefer cGMP over cAMP as a substrate (PDE1, PDE3 and PDE10)^59–61^. Intriguingly, CaMKK2-depletion resulted in a quantitative decrease in PDE1A protein levels, while not effecting other PDEs, in all three CaMKK2-KO clones of MDA-MB-231 cells (**Figure 7A, Supplementary Figure 4A**). Notably, of the three PDE1 isozymes, PDE1A has the highest affinity for cGMP, with a much lower Km for cGMP (5 μM) than for cAMP (112 μM)^62^. Pharmacological inhibition of CaMKK2 activity resulted in a similar reduction in PDE1A expression in both MDA-MB-231 cells and BT-20 cells (**Figures 7B, C**). Evaluation of the impact of CaMMK2 on *PDE1A* mRNA expression suggested that this regulatory activity was manifest at the level of transcription (**Supplementary Figure 4B**). Considering these results we postulated that CaMKK2 inhibits PKG1 activity and downstream phosphorylation of VASP by driving the expression of PDE1A, a key negative modulator of PKG1 activity. We took a pharmacological approach to test this hypothesis using vinpocetine, a PDE1-specific inhibitor^63^, IBMx, a pan-PDE inhibitor^64^, and sildenafil, a PDE5-specific inhibitor^65^. While neither IBMx nor sildenafil increased VASP phosphorylation, the PDE1-specific inhibitor, vinpocetine, increased serine 239 phosphorylation of VASP in parental MDA-MB-231 cells, phenocopying that which occurred in CaMKK2-depleted/inhibited cells (**Figure 7D**). Furthermore, inhibition of PDE1 increased phosphorylation of VASP in CaMKK2-WT cells exposed to ionomycin and, as expected, did not change the VASP phosphorylation levels in CaMKK2-KO cells (**Figure 7E**). Finally, inhibition of PDE1 with vinpocetine also reduced the migratory capacity of parental MDA-MB-231 (**Figure 7F, Supplementary Figure 4C**) and BT-20 cells (**Supplementary Figure 4D**).

**Figure 7.**
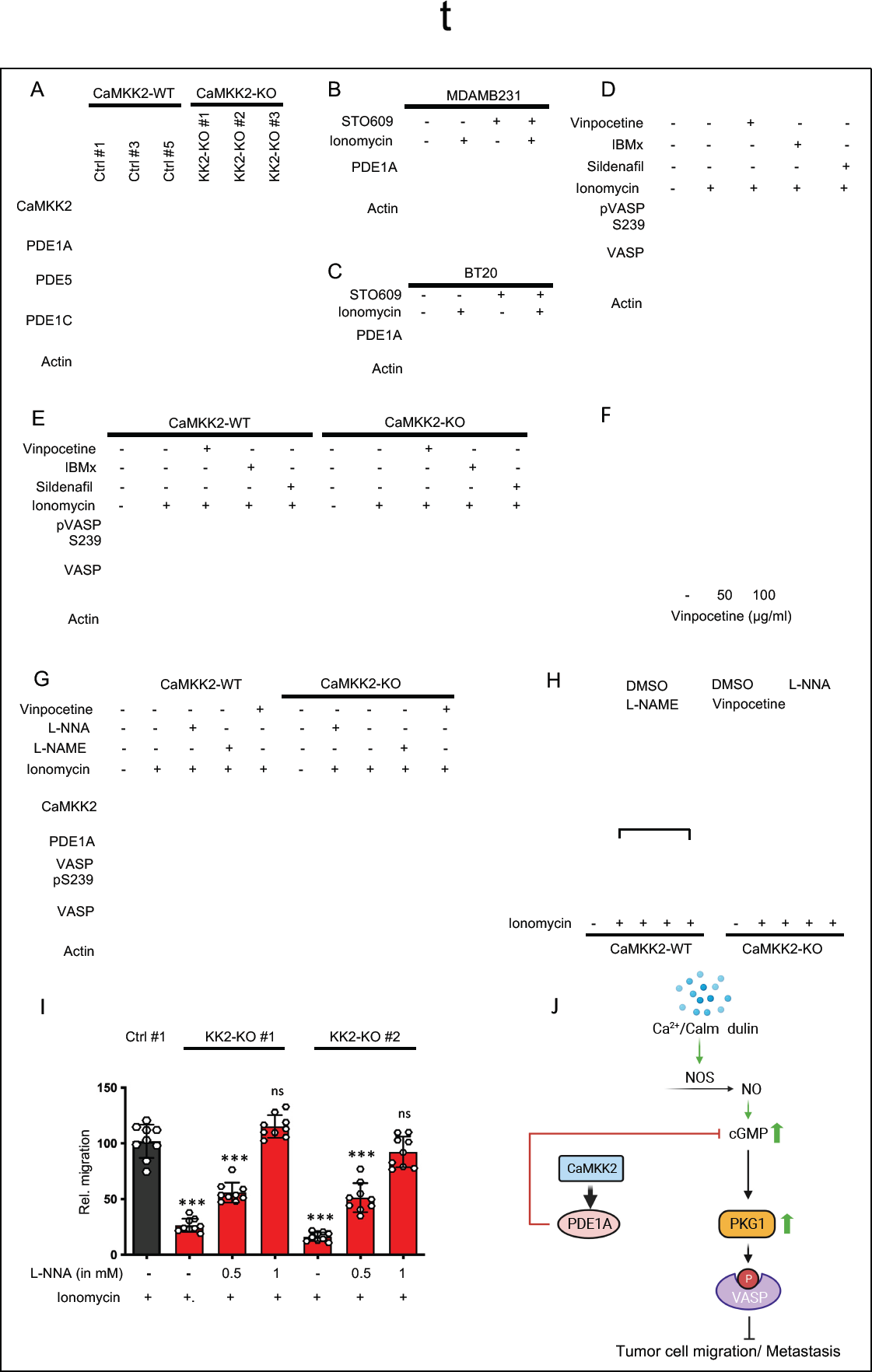
Depletion of CaMKK2 causes reduced expression of PDE1A, an upstream negative regulator of PKG1. **A,** Representative blot showing that depletion of CaMKK2 leads to significant decrease in PDE1A expression in MDA-MB-231 cells. Western immunoblots were used to analyze CaMKK2, PDE1A, PDE5 and PDE1C expression levels. β-Actin was used as a loading control. Representative results from four independent experiments are shown. **B, C,** Pharmacological inhibition of CaMKK2 (STO-609, 10µM) reduced PDE1A expression in MDA-MB-231 cells and BT20 cells. Representative blots from three independent experiments (**B**) and two independent experiments (**C**) are shown, **D,** Pharmacological inhibition of PDE1 with vinpocetine leads to increased phosphorylation of VASP at the Serine 239 residue in parental MDA-MB-231 cells. Vinpocetine (50µg/ml, 16h), IBMx – (100µM, 16h), sildenafil - (10µM,1h). MDA-MB-231 cells were pretreated with the inhibitors, as indicated, before exposure to either DMSO or ionomycin (1µM) with CaCL_2_ (1mM) for 2 hours. Western immunoblots to analyze phospho-VASP expression levels as compared to total VASP expression. β-Actin was used as a loading control. Representative results from four independent experiments are shown. **E**, Pharmacological inhibition of PDE1 with vinpocetine pretreatment leads to increased phosphorylation of VASP at the Serine 239 residue in CaMKK2-WT cells, while there is no change in CaMKK2-KO cells. Representative results from two independent experiments are shown. **F**, Pharmacological inhibition of PDE1 impairs the migratory ability of parental MDA-MB-231 cells in a dose-dependent manner. MDA-MB-231 cells were pretreated overnight with vinpocetine at increasing doses, as indicated, before exposure to ionomycin (1µM) with CaCL_2_ (1mM) for 2 hours. Cells were then harvested and allowed to migrate for 9 hours. Data are plotted as mean ± SEM; *n* = 4-5 random fields measurements from two individual experiments. **G, H,** Representative blot showing l-NNA (eNOS inhibitor) can block VASP phosphorylation of Serine 239 in CaMKK2-KO cells. CaMKK2-WT and CaMKK2-KO cells were cultured with NOS inhibitors (16h), as indicated, before being exposed to either DMSO or ionomycin (1µM) with CaCL_2_ (1mM) for 2 hours. Cell lysates were harvested and western immunoblots were performed. Representative blots from two independent experiments (**G**) and corresponding quantitative results for the band intensities of phospho-VASP vs. total VASP are shown (**H**). Data are plotted as mean ± SEM from two independent experiments. **I,** Pretreatment of CaMKK2-KO cell lines with L-NNA rescued the migratory phenotype in a dose-dependent manner. CaMKK2-WT and CaMKK2-KO cells were cultured in serum-starved media overnight and with or without L-NNA (1mM,16h), as indicated, before being exposed to ionomycin (1µM) with CaCL_2_ (1mM) for 2 hours. Cells were then harvested and allowed to migrate for 9 hours. Data are plotted as mean ± SEM; *n* = 4-5 random fields measurements from two individual experiments. Significance calculated relative to Ctrl#1. **J**, Schematic model of the mechanism by which CaMKK2 drives cell migration and metastatic dissemination in primary tumor cells. *P < 0.05, **P < 0.01, ***P < 0.005, *P* values were calculated using unpaired Student’s *t* test.

In this study we also observed that ionomycin-induced Ca^2+^ influx dramatically increased the inhibitory phosphorylation of VASP at the Serine 239 site. This was unrelated to effects on CaMKK2 as it also occurred in CaMKK2 depleted cells. Thus, we sought to understand how elevated calcium increased VASP phosphorylation secondary to activation of the cGMP/PKG1 axis. We hypothesized that this may relate to increased activity of soluble guanylyl cyclase (sGC) which synthesizes cGMP in response to increases in nitrous oxide (NO). This signaling molecule is synthesized by the nitric oxide synthases (NOS) of which there are three isoforms (NOS-1, NOS-2 and NOS-3^66^. NOS-1 (neuronal NOS) and NOS-3 (endothelial NOS) are constitutively expressed in most cells and require Ca^2+^ for activity, while NOS-2 (inducible NOS) activity is triggered by inflammatory stimuli, independent of Ca^2+^ influx^67 68^. Thus, we postulated that increased Ca^2+^ influx activates NOS-1 or NOS-3 (or both), thereby inducing cGMP production. Indeed, Nω-Nitro-L-arginine (L-NNA), a competitive inhibitor of NOS-1 and NOS-3, blocked ionomycin-induced VASP phosphorylation in both CaMKK2-KO and control cells (**Figures 7G, H**). This indicates that intracellular Ca^2+^ plays a key role in driving PKG1 activation/VASP phosphorylation by mediating cGMP production through the activation of NOS-1 and NOS-3. However, this effect is counteracted by CaMKK2/PDE1 activity to ultimately restrict aberrant PKG1 activity and downstream VASP phosphorylation in migratory cells. Of note, the negative impact of CaMKK2 knockdown on cell migration was reversed by treatment with L-NNA (**Figure 7I**). Taken together, our results indicate that CaMKK2 is required to maintain PDE1A expression in TNBC cells, which, by hydrolyzing cGMP attenuates PKG1 activity, to ultimately support tumor cell migration and spontaneous metastatic progression in breast cancer (**Figure 7J**).

### CaMKK2 expression is associated with poor progression-free survival and higher metastasis in high-grade serous ovarian cancer

We next addressed whether CaMKK2 expression was elevated and/or its activity associated with increased metastatic capacity in cancers other than those of the breast. Notably, it has been demonstrated that high-grade serous ovarian carcinoma (HGSOC) and TNBC are genetically and molecularly similar cancers, and share many underlying molecular features, such as *BRCA1* and *BRCA2* germline mutations and *TP53* somatic mutations^3^. Indeed, several ongoing clinical trials are designed to enroll patients with either HGSOC or TNBC, on the basis of the shared genetic characteristics of these two poor-prognosis cancers (ClinicalTrials.gov identifier: NCT01623349, ClinicalTrials.gov identifier: NCT00679783). As a starting place for our studies we used immunohistochemistry to evaluate CaMKK2 expression in tumors from 62 patients diagnosed with advanced stage HGSOC, fallopian tube or primary peritoneal cancer and found that higher expression (Intensity score >3) (**Figure 8A**) was directly correlated with worse progression-free survival (PFS) (**Figure 8B**) and overall survival (OS) (**Figure 8C**) in patients with advanced stage (Stage III) HGSOC. In a separate study of samples from patients with stage II to IV HGSOC, it was found that low expression of CaMKK2 (intensity score <3) was associated with longer PFS (**Figure 8D**) and a trend toward higher median OS in this expanded cohort was observed (**Figure 8E**). Our findings suggest that much like in TNBC, higher intratumoral CaMKK2 expression is predictive of highly aggressive phenotypes in patients with advanced stage HGSOC. Next, we carried out *in vivo* experiments to directly assess the impact of CaMKK2 inhibition on spontaneous metastasis in animal models of this disease. We found that pharmacological inhibition of CaMKK2 in animals injected with SKOV3ip cells, a validated model of HGSOC, resulted in significantly diminished spontaneous metastatic outgrowth, compared to vehicle-treated animals (**Figures 8F, G**), indicating that intratumoral CaMKK2 activity may be required to maintain metastatic capacity in HGSOC. From a clinical perspective, patients with HGSOC often present with metastatic disease, especially in the abdominal cavity. Despite standard-of-care cytoreductive surgery to resect tumor burden from any involved organs in the abdominal and pelvic cavity, most HGSOC patients remain at high risk of recurrence following primary treatment due to secondary metastasis beyond the abdominal cavity. It was significant, therefore, that in addition to impacting overall metastatic burden, inhibition of CaMKK2 activity significantly reduced invasive metastatic outgrowth in secondary sites, distant from the abdominal and pelvic cavity (**Figure 8H**). Our results suggest that targeting CaMKK2 may be an effective therapeutic approach to limit secondary relapse in HGSOC. These findings highlight the potential translational benefit of utilizing CaMKK2 inhibitors to limit long-term recurrence in high-risk cancers like TNBC and HGSOC.

**Figure 8.**
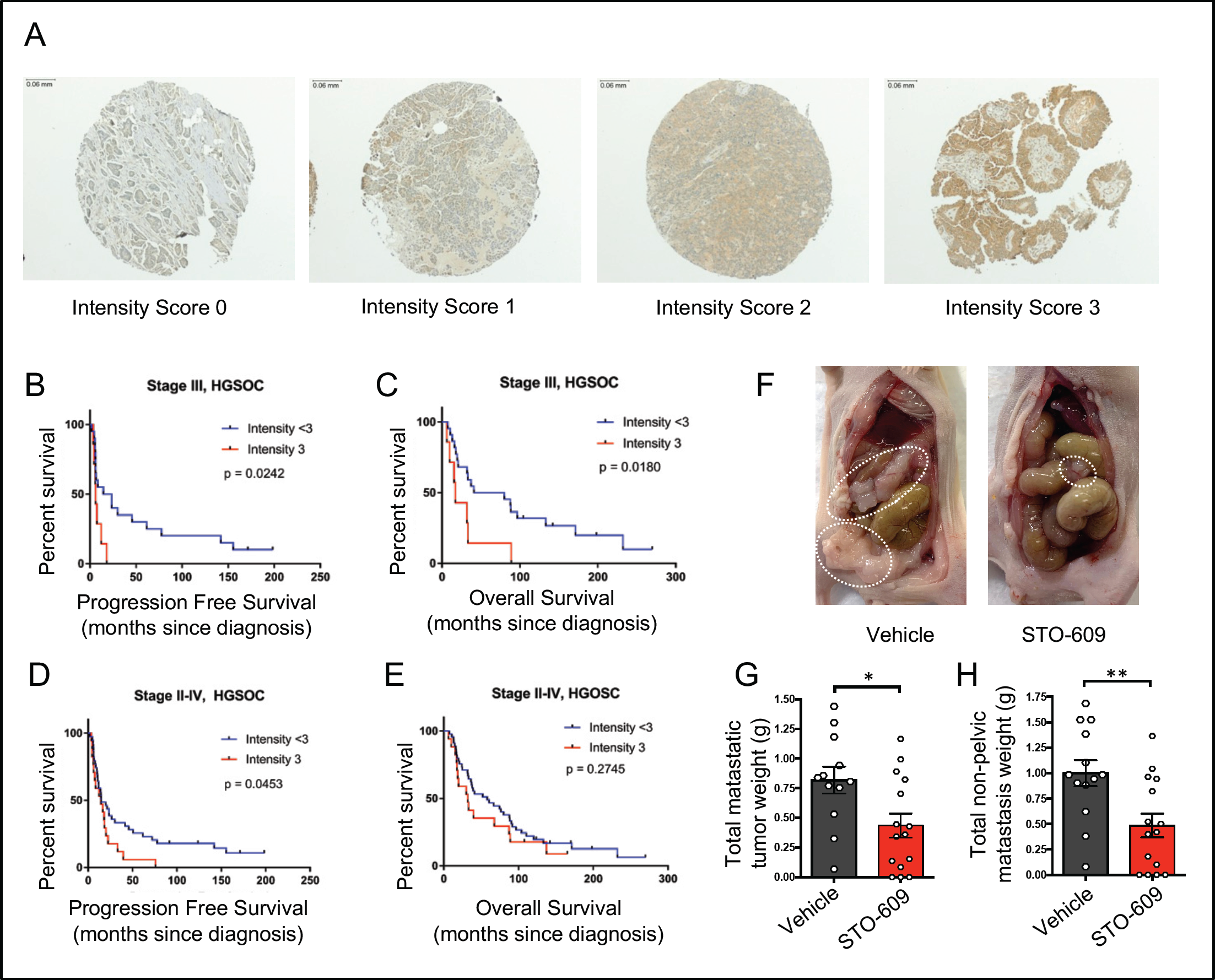
CaMKK2 expression is associated with lower survival rates in patients with highgrade serous ovarian cancer (HGSOC) **A,** Representative photomicrographs of tumor microarray sections stained for CaMKK2 by immunohistochemistry with corresponding intensity scores (10x magnification). **B,** Progression-free survival (PFS) and **C**, overall survival (OS) of patients with stage III HGSOC were evaluated and demonstrated that low CaMKK2 expression (intensity score <3) was associated with a longer median PFS (19.1 months vs 6.2 months, p=0.0242) and OS (60.6 vs 17.1 months, p=0.0180) than those with high expression. **D**, PFS of patients with stage II to IV HGSOC were also evaluated, which demonstrated that low CaMKK2 expression was associated with a longer median PFS than patients with high expression (14.7 months vs. 14.5 months, p=0.0453) in this expanded cohort. **E**, OS of patients with stage II to IV were similarly evaluated and showed a trend toward higher median OS in patients with low expression (58.0 months vs. 32 months, p=0.2745). **F**, 8-week-old female nude mice were inoculated with SKOV3ip1 cells (intraperitoneally [IP]) and received vehicle (control) or STO-609. Representative images. **G**, weight of metastatic tumor burden and **H**, weight of non-pelvic metastasis are shown (n=15 per group). Data are expressed as the mean ± SEM. \**P <* 0.05, \*\**P <* 0.01, \*\*\**P <* 0.005, and \*\*\*\**P <* 0.001, by Student’s *t* test (**G** and **H**) and logrank test (**B, C, D, E**).

## Discussion

The development of metastasis requires tumor cells to dissociate from their primary site, acquire migratory capability and invade the basement membrane and surrounding tissue to ultimately gain vascular access^69–71^. Only a small percentage of cells that are shed from the primary tumor possess these invasive (and metastasis-initiating) properties. Here we show that tumor cell intrinsic expression of CaMKK2 is a key determinant of the metastatic potential of TNBC cells. Specifically, genetic ablation of CaMKK2, or its pharmacological inhibition, dramatically reduced the invasiveness and migratory potential of cell lines modeling TNBC *in vitro* and significantly reduced spontaneous metastatic outgrowth from primary tumors *in vivo*. Importantly, sustained CaMKK2 activity was also found to be required to maintain metastatic capacity in HGSOC, a high-risk, poor prognosis ovarian cancer subtype that shares many genetic features with TNBC. Mechanistically, our results implicate an intricate signaling axis in which CaMKK2 plays a central role in controlling intracellular actin dynamics to favor heightened tumor cell motility and migratory capability. We find that CaMKK2 restricts PKG1 activity in metastatic tumor cells by driving the expression of PDE1A, a key phosphodiesterase that catalyzes the degradation of intracellular cGMP. This implicates CaMKK2 as a key driver of the negative regulatory mechanism that counteracts NOS-mediated cGMP synthesis to ultimately maintain low levels of PKG1 activation and limit the inhibitory phosphorylation of VASP in metastatic tumor cells. Loss of CaMKK2 shifts the balance towards increased intracellular cGMP production, thereby leading to enhanced PKG1 activation and increased phosphorylation of VASP at the Serine 239 site. This inhibitory phosphorylation event impairs VASP-driven actin polymerization activities and stress fiber assembly to ultimately impact tumor cell motility and metastatic capability. Our data highlights the potential clinical utility of targeting CaMKK2 as a strategy to diminish the invasiveness and metastatic potential of both breast and ovarian cancers.

Previously, we reported that CaMKK2 is expressed within myeloid and epithelial cells in all human breast tumor subtypes and that its overexpression is associated with aggressive triple-negative disease and poorer disease outcomes^31^. Our updated analysis further demonstrated that its overexpression is associated with poorer disease outcomes in TNBC and HGSOC. To our knowledge, this is the first report which demonstrates that intratumoral CaMKK2 plays a central role in regulating metastatic capacity in breast and ovarian cancers. Although CaMKK2 has been demonstrated to have distinct pro-tumorigenic roles in several cancers^28 30 72^, the mechanisms by which it impacts tumor metastasis have not been elucidated. A cell-based study implicated CaMKK2 activity in medulloblastoma migration^73^, and we previously reported that CaMKK2 inhibition impaired androgen-dependent invasion/migration of prostate cancer cells *in vitro*^27^. More recently, CaMKK2 activity was shown to decrease anoikis, thereby aiding metastatic progression in lung cancer^22^. These observations, and our new data showing that elevated CaMKK2 expression is associated with aggressive disease phenotypes in both TNBC and HGSOC, highlights the importance of this enzyme in tumor pathobiology and suggests that it may be a useful therapeutic target for the clinical management of a number of different tumor types.

VASP is a substrate of cGMP-dependent protein kinases (PKA and PKG) which localizes to regions of dynamic actin remodeling to promote F-actin assembly where it integrates signaling events that regulate cytoskeletal reorganization and cell movement^46 53 74–76^. Originally, VASP phosphorylation was described in platelets, where both PKA and PKG were shown to phosphorylate the residues Serine 157 and Serine 239 respectively^77^. Notably, the S239 site is adjacent to the VASP G-actin binding site and phosphorylation at this residue inhibits its ability to bind actin causing a significant reduction in VASP-driven F-actin accumulation^53^. It was significant therefore, that CaMKK2 deficiency increased S239 phosphorylation of VASP leading to disrupted stress fiber assembly in TNBC cells, an activity that dramatically impaired their motility and migratory capability. Our results indicate that sustained CaMKK2 expression drives tumor cell motility by negatively regulating PKG1 activity in migratory cells, thereby limiting the inhibitory phosphorylation of VASP at the S239 site. Apart from facilitating stress fiber assembly within migratory cells, VASP-dependent actin polymerization is crucial for the formation of actin-containing protrusions (filopodia and lamellipodia) and invadopodia that enable cells to invade through the basement membrane^78–81^. Future studies will explore the direct connections between CaMKK2, VASP and the formation of these specialized structures.

The NO/cGMP/PKG1 pathway is involved in the regulation of several different processes in cells and is tightly regulated by phosphodiesterase-dependent hydrolysis of cGMP^59 82 83^. It is significant, therefore, that CaMKK2 was found to be an upstream regulator of PDE1A expression and that depletion of CaMKK2 induced the activity of PKG1, a major downstream effector of the NO/cGMP signaling axis. Notably, VASP is a primary downstream substrate for PKG1 and the phosphorylation of this protein at the S239 site is used as a readout for PKG1 activity^84–86^. Our data suggest that VASP is the primary biochemical link between PKG1 and increased cell motility/metastasis. However, a recent report demonstrated that PKG1 can also phosphorylate TSC2 to inhibit mTORC1 signaling^87^. The role of this pathway in cancer pathobiology requires further investigation.

We have shown that the activity of the NO/cGMP/PKG1 pathway, as measured by VASP phosphorylation, is upregulated significantly upon elevation of intracellular calcium. Whereas CaMKK2 itself can be activated in response to elevated intracellular Ca^2+^, several studies have demonstrated that this enzyme has significant activity independent of Ca^2+^ signaling (autonomous activity)^10 88^. Indeed, we demonstrated that CaMKK2-dependent increases in PDE1A expression in the cell lines tested were unaffected by the addition of a calcium ionophore indicating that the effect of this enzyme on cellular migration through the PDE1A/cGMP/PKG1 signaling axis is a function of its Ca^2+^-independent autonomous activity. Interestingly, PDE1 is the only member of the phosphodiesterase family that is activated by Ca^2+^/CaM binding^89^. Indeed, in conditions of high calcium, PDE1A replaces PDE5 as the primary phosphodiesterase responsible for the hydrolysis of intracellular cGMP^90^. Thus, it appears that while the entry of Ca^2+^ induces cGMP production through the NO/cGMP/PKG1 signaling axis, it also initiates a negative feedback mechanism that results in PDE1A-induced cGMP degradation. Loss of CaMKK2, and subsequently decreased expression of PDE1A, results in elevated PKG1 activity. Defining the specific mechanisms by which CaMKK2 regulates PDE1A expression (and activity) is a current research priority.

The somatic molecular profiles of TNBC and HGSOC share many similarities, including frequent *TP53* mutations, inactivation of *BRCA1* and *BRCA2,* mutations in *NF1* and *RB1* and overall poor prognosis^3 91^. Patients with HGSOC often present at an advanced stage (Stage III) with metastatic disease, especially, in the pelvic and abdominal cavity. The standard of care for these patients is cytoreductive surgery to remove tumors in the abdominal region, followed by adjuvant chemotherapy. Following primary treatment, most patients with HGSOC have a complete clinical, radiographic and biochemical response. However, these patients usually develop secondary metastasis within 1-2 years of cytoreductive surgery, despite the adoption of approved maintenance therapies that include poly ADP ribose polymerase inhibitors^92–96^. Given that the five-year overall survival for patients with Stage III and IV disease remains ∼ 20-40%^97^, improving upon the current standard of care in HGSOC represents a critical unmet need. Our data which demonstrated a very strong protective effect of CaMKK2 inhibitors in animal models of advanced ovarian cancer highlight a potential role for these drugs in the long-term management of this disease by preventing/slowing down the metastatic progression of post-operative residual disease. Similarly, TNBC is the most aggressive subtype of breast cancer and the tumors tend to become highly invasive early during cancer development. Recent data have suggested that isolated local or regional relapses (ILRRs), defined as clinically documented relapse in the ipsilateral breast or regional nodes within one year of the primary diagnosis, significantly increased the risk of distant recurrence and death in breast cancer patients^98^. This suggests that TNBC patients who present with lymph node-positive, early-stage disease or develop local relapses within a year of first-line treatment may benefit from neoadjuvant combination therapies targeting CaMKK2 in conjunction with standard chemotherapy.

The immune checkpoint inhibitor pembrolizumab (anti-programmed cell death protein-1 (PD-1) monoclonal antibody) was recently approved for use in high-risk, early-stage TNBC, in combination with neoadjuvant chemotherapy and as a single agent in the adjuvant setting post-surgery^99^. Further, there are ongoing clinical trials evaluating the effectiveness of pembrolizumab in platinum-resistant advanced ovarian cancer (ClinicalTrials.gov Identifier: NCT02440425). It is significant therefore that we demonstrated in a previously published study that targeting myeloid-specific CaMKK2 in the mammary tumor microenvironment significantly enhances anti-tumor immune responses by reducing the immunosuppressive activity of tumor associated macrophages and increasing the recruitment of cytotoxic T cells into the tumor^31^. This suggests that CaMKK2 inhibitors may increase the efficacy of immune checkpoint inhibitors in cancers that are enriched in tumor associated macrophages. The observation in this study that CaMKK2 also regulates the metastatic potential of cancer cells suggests that CaMKK2 inhibitors will have a dual action in tumors and when used with ICBs could significantly advance the pharmacotherapy of breast and ovarian cancers.

## Materials and Methods

### Reagents

STO-609 and GSK1901320 (referred to as GSKi) were synthesized at the Duke University Small Molecule Synthesis Facility. Ionomycin (Catalog # 10004974) was obtained from Cayman Chemicals and solubilized in DMSO at room temperature, sterile filtered (0.2 µM), and stored at −20 °C. The pan-PDE inhibitor, IBMX (Catalog # BML-PD140) and PDE1-specific inhibitor, vinpocetine (Catalog # BML-PD185) were purchased from Enzo Life Sciences. Sildenafil (Catalog # 10008671) was obtained from Cayman Chemicals. The nitric oxide synthase (NOS) inhibitors, Nω-Nitro-L-arginine (L-NNA) (Catalog # N5501) and Nω-Nitro-L-arginine methyl ester hydrochloride (L-NAME) (Catalog # N5751) were both obtained from Millipore Sigma and solubilized in 1M HCL and dH2O, respectively. To stain filamentous actin, rhodamine-phalloidin (Catalog # R415) was purchased from Thermo Fisher Scientific, dissolved in methanol and stored at -20°C. The small molecule PKG inhibitors, RKRARKE (Catalog # 370654) and KT5823 (Catalog # K1388) were both obtained from Millipore Sigma.

### Cells and Cell culture conditions

Luciferase-labeled lung-trophic MDA-MB-231-4175 (here referred to as MDA-MB-231) cells were kindly provided by Dr. Joan Massague (Memorial Sloan Kettering Cancer Center). BT-20 and HCC1954 cells were purchased from American Type Culture Collection (ATCC), Manassas, VA. SKOV3ip1 and HOC7 cells were kindly provided by Drs. Guillermo Armaiz-Pena (Ponce Health Sciences University) and Arktak Tovmasyan (Duke University), respectively. MDA-MB-231 and BT20 cells were maintained in Dulbecco’s Modified Eagle’s Medium (DMEM). HCC1954, HOC7 and SKOV3iP1 were grown in RPMI-1640 media. DMEM and RPMI media were supplemented with 8% fetal bovine serum (FBS, Sigma), 1 mM sodium pyruvate, and 0.1 mM nonessential amino acids (Thermo Fisher Scientific). All cells were cultured in a humidified incubator at 37 °C with 5% CO2. For experiments with ionomycin, cells were grown till 90% confluent and the culture media was then replaced with Hanks’ Balanced Salt Solution (HBSS) (Gibco) supplemented MgCL2 (1mM) for 15 minutes prior to ionomycin exposure. The cells were then treated with either DMSO or ionomycin (1µM) with CaCL2 (1mM) for varying time periods (as indicated) before harvesting the cells for downstream processing.

### Generation of CaMKK2 knockout and addback cell lines

sgRNAs targeting specific sequences at the N-terminal end of the *CaMKK2* gene were purchased from Synthego Corp., Menlo Park, CA. For each sgRNA, a ribonucleoprotein (RNP) complex was formed with the Cas9 2NLS nuclease (Synthego), at a sgRNA: Cas9 ratio of 6:1. MDA-MB-231 cells were seeded at 8 × 10^4^ cells/ml and transfected with the RNP complex using Lipofectamine CRISPRMAX transfection reagent (Catalog # CMAX00008, Thermo Fisher Scientific). 5-6 days post transfection, cells were plated for single cell isolation in 96 well plates at a limiting dilution of 0.6 cells/well. Single cell clones were allowed to expand and clones with successful knockout of CaMKK2 was identified directly by immunoblotting. One random clone was selected for each of sgRNA #1, #2 and #3, as indicated, and used for experiments. The sequences of all sgRNAs used in the study are included in **Supplementary Table I**.

Lentiviral constructs of wildtype CaMKK2 were created by shuttling CaMKK2 from the pDONR221 vector backbone to pLenti-CMV-puro-DEST (Addgene) using Gateway cloning. All plasmids were confirmed via restriction digests and/or Sanger sequencing. CaMKK2 addback cell lines were generated by infecting the MDA-MB-231 CaMKK2 knockout clones (KK2-KO #1 and KK2-KO #2) with lentivirus expressing wildtype CaMKK2. Transduced cells were selected for using puromycin treatment (1 µg/mL). Addback was confirmed by immunoblot analysis.

### siRNA transfection

MDA-MB-231 (50,000 cells/ml), BT-20 (50,000 cells/ml), HCC1954 (60,000 cells/ml) and HOC7 (75,000 cells/ml) cells were transfected with either control siRNAs (Catalog # 12935146, Thermo Fisher Scientific; Catalog # SIC001, Millipore Sigma) or CaMKK2 siRNAs (50nM) (Catalog # 10620318, Thermo Fisher Scientific) using the DharmaFECT 1 transfection agent (Catalog # T-2001-03, Dharmacon) according to the manufacturer’s instruction. Cells were collected for downstream analysis after 48 hrs (RNA) or 72 hours post transfection (protein). The sequences of the siRNAs used are included in **Supplementary Table II**.

### Migration/invasion assays

Migration of cancer cells was evaluated using transwell migration and wound healing assays. Cell invasiveness was tested by utilizing invasion assay chambers. For transwell migration, cells were cultured in serum-starved conditions overnight and 5 × 10^4^ cells were seeded in the upper insert of a Transwell chamber (8 µm pore size polycarbonate membrane filter, Catalog # 3422, Corning) in 0.25 ml of 0.5% FBS-containing media, with 0.8 ml of 8% FBS-containing media in the lower chamber. Cells were incubated for 3–12 hrs as indicated in individual experiments. To evaluate invasiveness, cells were seeded on Biocoat invasion chambers with 8 µm pore size PET membrane precoated with extracellular matrix proteins (Catalog # 354480, Corning). Invasion assays were conducted as per the manufacturer’s protocol using the same seeding conditions as in transwell migration. For both migration and invasion assays, cells that migrated through the membrane were fixed with 4% paraformaldehyde and stained with 0.1% crystal violet for 15 min at room temperature and photographed under the microscope. A minimum of nine images were photographed, and each sample condition was performed with at least three technical replicates. Total number of migrated/invaded cells were measured by Image J. For migration/invasion experiments with CaMKK2 inhibitors, cells were first treated with Vehicle or STO-609 (10 µM) or GSKi (1 µM) in 8% FBS-containing media for 24 hours and then again in serum-starved media for the next 24 hours before the cells were harvested for experiments (total treatment time: 48h). For wound healing assays, cells were seeded in a 12-well plate and cultured to a confluent monolayer, media was then replaced with serum-free (or 0.1% FBS-containing media) and cultured overnight. The next day, two scratches were introduced per well by scraping the monolayer with a p200 pipette tip and marked for orientation. At least twelve images were photographed by microscopy immediately (0h) and after 48h and 60h. Scratch introduced open area (OA) was measured by Image J and evaluated by Migration Index = [(OAT1 − OAT2)/OAT1] × [OAT1/Ave of all OAT1].

### Animal Studies

6-week old Athymic nude-*Foxn1^nu^* mice were purchased from Envigo. The mice were housed in secure animal facility cages under pathogen-free conditions on a 12-hour light/12-hour dark cycle at a temperature of approximately 25°C and 70% humidity. Mice had ad libitum access to food and water. All xenograft procedures were approved by the Duke University Institutional Animal Care and Use Committee (IACUC). To study spontaneous metastasis from the primary tumor, luciferase-labeled cells (1×10^6^ cells per mouse) (suspended in HBSS/Matrigel, 1:1) were inoculated orthotopically into the mammary fat pads of 8-week old female athymic nude mice. Mice were monitored daily, and tumor volume was measured with electronic calipers using the formula V=Lx(WxW)/2. Each mouse in each group was euthanized when the tumor reached a maximum size of 2000-2500 mm^3^ as specified by the Duke IACUC protocol. Prior to euthanizing, the mice were injected with D-luciferin. The secondary organs (liver, lungs) were then collected and micrometastasis was visualized by bioluminescent imaging *ex vivo* using the IVIS Lumina XR In Vivo Imaging System (Perkin Elmer). Experimental metastasis was studied by injecting viable luciferase-labeled metastatic cells (500,000) (suspended in ice-cold 200 µl HBSS) into the lateral tail veins of 8-week old athymic nude mice. The extent of lung metastasis was monitored by bioluminescent imaging of the live animals using the IVIS Lumina XR In Vivo Imaging System (Perkin Elmer). Mice were euthanized at indicated time points. For *in vivo* experiments involving drug treatments, 8-week old athymic nude mice were orthotopically injected with parental MDA-MB-231 cells (1.5×10^6^ cells per mouse) into the mammary fat pad, as described above. Therapeutic dosing was started 5 days after tumor cell injection at a starting tumor size of 50mm^3^ approx. The mice were dosed with Vehicle (PEG-8, PolyethyleneGylcol 400; Spectrum Chemical MFG Corp, Gardena, CA), STO-609 (40µmol/kg body weight) or GSKi (10µmol/kg body weight) via intraperitoneal injections every third day. Tumor size was monitored throughout the study and micrometastasis in secondary organs (lung, liver) was evaluated as described above. For *in vivo* metastasis experiments with the high grade serous ovarian cancer model, SKOV3ip1 cells (1 × 10^6^ cells in 200 μl of HBSS) cells were injected intraperitoneally into 9-week old female athymic nude mice (Charles River, Wilmington, MA). Mice were treated via intraperitoneal injection with vehicle (PEG-8, PolyethyleneGylcol 400; Spectrum Chemical MFG Corp, Gardena, CA) or STO-609 (MedKoo Biosciences, Inc., Morrisville, NC; 40 µmol/kg) twice weekly. Once mice in any group became moribund, all mice were sacrificed and necropsied. Metastatic burden was harvested, weighed, and location of metastatic nodules were recorded. Statistical analyses were performed by Student’s t-test or two-way ANOVA followed by a Bonferroni multiple comparison test.

### Sandwiched White Adipose Tissue (SWAT) preparation

Control cells (Ctrl#1, Ctrl#3) and CaMKK2-KO cells (KK-KO#1, KK2-KO#2) were stained with Cell Brite Green Dye at 37°C for 45 minutes and mixed with freshly isolated minced human breast tissue in individual 3.5 cm SWAT petri-dish as described^39 40^. The SWAT dishes were incubated overnight to allow the system to stabilize. Media was changed after 24 hours and the SWAT systems received fresh media daily till the end of the experiment.

### Time-lapse Video and measurement of 3D cell movement

Using the Nikon Elements Software, a workflow was created to capture fluorescence microscopy and bright field images every 5 minutes over 6 hours yielding a total of 73 frames. Each sample (both control lines and both CaMKK2 knockout lines) were captured using fluorescence only, followed by merged fluorescence and bright field images. Each SWAT petri dish was then transferred sterilely to the Nikon scope top incubator and fixed in place. The incubator was set to 37°C and 5% CO2 for all videos. Using FITC and bright field lenses, specific areas of interest were selected that included both easily identifiable adipocytes and several of the dyed cancer cells. Magnification was adjusted to 20X and auto exposure for each field type was adjusted to provide better picture quality. Each SWAT sample was filmed for a total of 12 hours and the resulting videos were given scale bars, exported, and compressed to AVI format. Each video is approx. 14 seconds long (5 fps). AVI files were opened using the ImageJ software and the tracking plugin MTrackJ was loaded through which specific cells were marked and traced through every frame of a given AVI file in order to generate a relative measurement of cell movement given in pixel length. Each track is represented by different colors and numbers for easy identification. Statistical analyses to evaluate differences in 3D movement of cells were performed by Student’s t-test.

### Immunoblotting and Quantitative PCR

Cells were washed three times with 2 ml of ice-cold PBS, snap-frozen and lysed with 0.15 ml of phospho-RIPA lysis buffer (Tris-HCl pH 7.5, 50mM; NaCl, 150 mM; NP-40, 1%; Sodium deoxycholate, 0.5%; SDS, 0.05%; EDTA, 5mM; Sodium fluoride, 50mM; Sodium pyrophosphate, 15mM; glycerophosphate, 10mM; Sodium orthovanadate, 1mM) with protease inhibitor cocktail (Millipore-Sigma, P-8340). 25µg of lysates were denatured and resolved by SDS-PAGE. Proteins were transferred to Odyssey Nitrocellulose Membrane (Catalog # 926-31092, LI-COR Biosciences). Primary antibodies used were anti-CaMKK2 (MO1, clone 1A11, 1:2000, Catalog # H00010645-M01, Abnova); anti-phospho AMPKα (Thr 172) (1:1000, Catalog # 2531, Cell Signaling); anti-AMPKα (1:500, Catalog # 2532, Cell Signaling); anti-phospho-VASP (Ser239) (1:1000, clone 16C2, Catalog # 05-611, Millipore Sigma); anti-phospho-VASP (Ser157) (1:1000, Catalog # 3111, Cell Signaling); anti-VASP (1:2000, Catalog # HPA005724, Atlas Antibodies); anti-PDE1A (1:500, Catalog # PD1A-101AP, FabGennix); anti-PDE5 (1:1000, Catalog #2395, Cell Signaling); anti-PDE1C, 1:2000, Catalog # ab14602, Abcam); anti-Actin (1:10000, 8H10D10, Catalog # 3700, Cell Signaling). Secondary antibodies used were HRP-conjugated anti-mouse IgG (1:5000) Catalog # 1706516 and anti-rabbit IgG (1:5000) Catalog # 1706515, BioRad) and protein bands were visualized by Western Lightning Plus ECL system (Catalog # ORT2655 and ORT2755, Perkin Elmer).

For quantitative PCR, RNA was isolated using RNA Aqueous Micro kit (Catalog # 1931, Ambion) followed by cDNA synthesis using an iScript cDNA synthesis kit (Catalog # 170-7691). Quantitative amplification was performed using Sybr Green (Catalog # 1725124, Bio-Rad) and a CFX-384 Real Time PCR detection system. Primers used to amplify the *PDE1A* gene are 5’-GAG TCT TCC CTC AGC TTT GT-3’ (forward) and 5’-TTT CAG TCT GTT CTC CTG TAA GAT-3’ (reverse).

### Kaplan Meier survival plot

To generate the Kaplan Meier plots (https://kmplot.com) in Figure 1, the parameters were set as follows -- the probe to “CaMKK2 210787_s_at”, the time to “(months): 180”, the cutoff to “Upper quartile”, the survival to “OS (overall survival)”, and the PAM50 Subtype to “Basal, Luminal A, Luminal B, and HER2+”. As of October 29, 2021, the number of patients included in KMplot analysis was 1879. The survival data were downloaded as text files and used to draw the survival plots using Survminer R package (version 0.4.9).

### Gene signature analysis

TCGA aggregate expression datasets for breast cancer were retrieved from NCI Genomic Data Commons and processed using TCGAbiolinks (version 2.16.4) and SummarizedExperiment (version 1.18.2) R packages. The samples’ definition and PAM50 subtype were downloaded alongside the expression datasets. Tumor samples without PAM50 annotation were excluded. The number of samples included in the downstream analysis by definition: 1072 primary solid tumor and 113 normal. Normal samples are designated as normal-adjacent. The number of samples by PAM50 subtyping: 189 Basal, 82 Her2, 206 LumB, 555 LumA, and 40 normal-like. EdgeR R package (version 3.30.3) was employed to normalize the RNAseq raw count for the library depth. A matrix using samples definition was designed, and the raw count and the normalization factor was calculated for normalization. Gene signature analysis was performed using the Gene Set Variation Analysis (GSVA) R package (BMC Bioinformatics, 14, 7. doi: 10.1186/1471-2105-14-7). The indicated gene sets were defined and the score in each sample was calculated. The method in the GSVA analysis were set to “gsva”, and the kcdf option to “Gaussian”. When comparing the scores of the gene sets between CAMKK2-high or CAMKK2-low expressing samples within different PAM50 subtypes, Wilcoxon signed-rank test was used to calculate the p values. The box plots and the scattered plots were drawn using ggplot2 (version 3.3.5).

### Analyses of human HGSOC correlates

We evaluated information from 62 patients diagnosed with advanced stage high-grade serous ovarian, fallopian tube or primary peritoneal cancer who were chemotherapy naive and with available bio-banked formalin-fixed paraffin-embedded (FFPE) blocks. Histologic diagnoses were confirmed by a board-certified anatomic pathologist with experience in gynecologic pathology (K.C.S). Clinico-pathological records were retrieved and stored in a database. The study was approved by the Institutional Review Board (IRB Pro00013710). After routine deparaffinization in xylene, rehydration, and inhibition of endogenous peroxidase activity with 3% hydrogen peroxidase, sections were exposed to heat-induced epitope retrieval (HIER) with the Decloaking Chamber (Biocare Medical, Pacheco, CA), in Citra buffer (pH 6.0). After blocking with Background Terminator (Biocare Medical, Pacheco, CA), the sections were incubated with rabbit anti-human antibody directed against CaMKK2 (HPA017389, Sigma, St. Louis, MO) at 1:500, overnight at 4 °C. This was followed by incubation with 4 Plus Universal Detection system (Biocare Medical, Pacheco, CA), a universal affinity purified biotinylated secondary antibody, which is bound to a horseradish peroxidase (HPR)-labeled streptavidin. The chromagen/substrate was applied to visualize the location and intensity of the protein of interest, and slides were counterstained with hematoxylin. Protein expression of CaMKK2 was evaluated using immunohistochemistry. Intensity of staining was reported using a semi-quantitative 4-part scale, as shown in the figure. Time-to-event analyses were based on the Kaplan-Meier method, and events were compared using the log-rank (Mantel-Cox) test.

### Statistics

Statistical analysis was performed with GraphPad Prism 8.0 (GraphPad Software), using either a 2-tailed Student’s *t* test or 1- or 2-way ANOVA. For both 1-way and 2-way ANOVAs, a post-test analysis was performed using Bonferroni’s multiple correction. The number of replicates is indicated in the figure legends. A *P* value of less than 0.05 was considered statistically significant.

### Study approval

All animal experiments were performed according to guidelines established and approved by the IACUC of Duke University

## Acknowledgments

**Funding:** This project was supported by US Department of Defense Award (BC191370) to D.P.M. R.A.P. is supported by grants from the NIH 1K12HD103083-01, AAOGF-GOG Foundation and the Emerson Collective. S.P. and J.F. are supported by NIH Basic Research in Cancer Health Disparities R01 Award (R01CA220314).

**Author contributions:** D.M., C.Y.C. and D.P.M. conceived of and designed all the experiments. D.M. performed the experiments with technical support provided by R.A.P., C.N.H., P.K.J., B.C., S.A., R.W. and C.Y.C. M.A.A. and J.A.F. contributed to Figure 1. K.L.H. and J.A.F. performed experiments using the SWAT preparation technique with human breast tissue which was overseen by F.H.L. and M.B. K.C.S. performed the analysis of human HGSOC correlates. R.A.P. contributed to Figure 8 and helped in data interpretation. Manuscript was written by D.M. and D.P.M. with critical inputs from C.Y.C. The project was managed and overseen by D.P.M.

**Data and materials availability:** All data are available in the main text or the supplementary materials

