## Supplementary Figures and Tables for "Ca^2+^/Calmodulin Dependent Protein Kinase Kinase-2 (CaMKK2) promotes Protein Kinase G (PKG)-dependent actin cytoskeletal assembly to increase tumor metastasis"

### CaMKK2 regulates actin cytoskeletal remodeling within tumor cells to facilitate tumor invasiveness and metastasis

**This PDF file includes:**

Supplementary Figures 1-4
Supplementary Videos 1-4 (legends)
Supplementary Tables 1-2

**Other Supplementary Materials for this manuscript include the following:**

Supplementary Videos 1-4

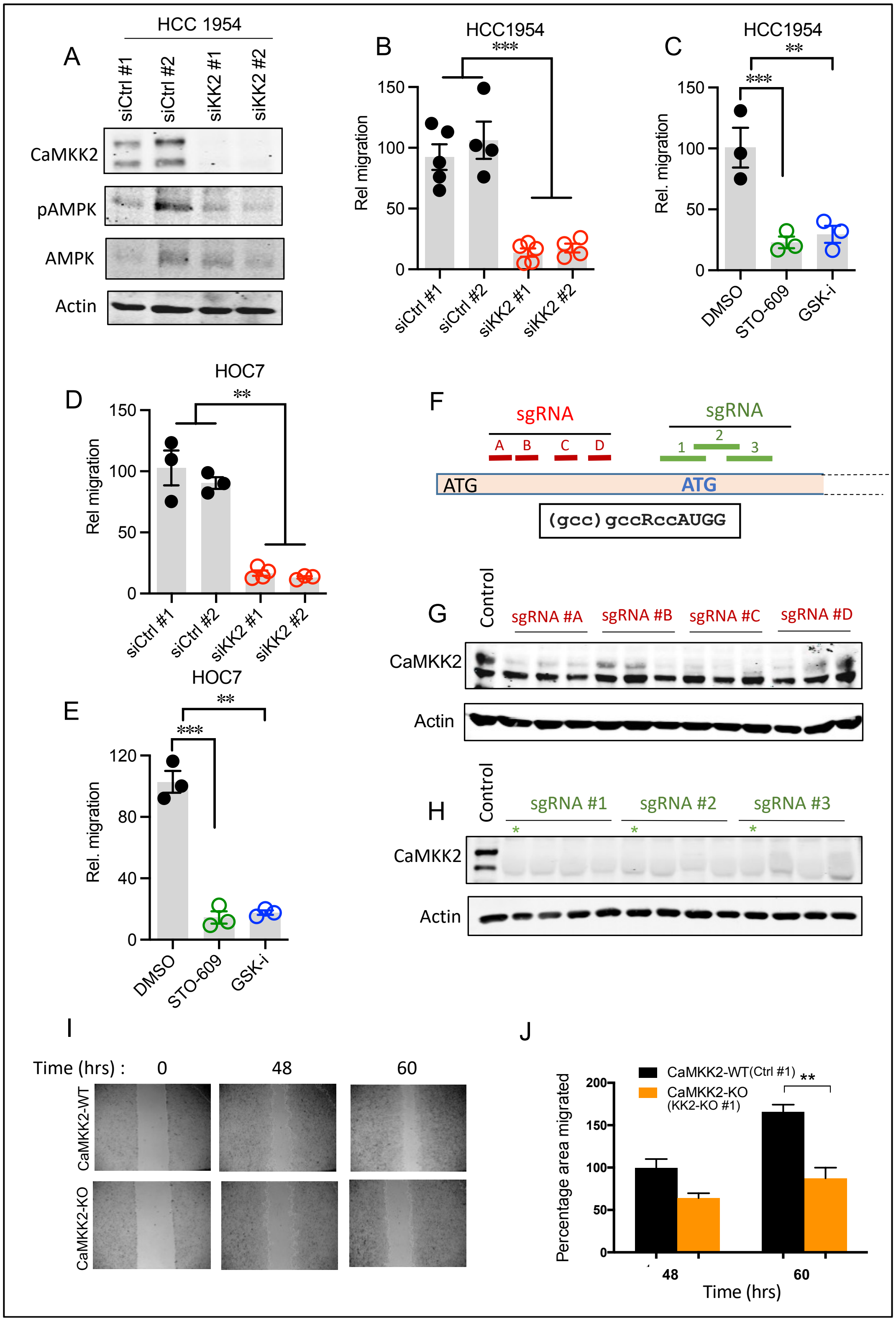


**Supplementary Figure 1. Ablation of CaMKK2 impairs migration in invasive breast cancer cells. Related to Figure 2. A,** Representative blots showing knockdown of CaMKK2 in HCC1954 breast cancer cells. **B,** CaMKK2 knockdown impaired migratory ability in HCC1954 cells *in vitro*. **C,** Pharmacological inhibition of CaMKK2 with STO-609 (10μM, 48h) and GSKi (1μM, 48h) resulted in impaired migration in HCC1954 cells *in vitro*. **D,** CaMKK2 knockdown impaired migratory ability in HOC7 ovarian cancer cells *in vitro.* **E,** Pharmacological inhibition of CaMKK2 with STO-609 (10μM, 48h) and GSKi (1μM, 48h) resulted in impaired migration in HOC7cells. **F,** Schematic representation of the CaMKK2 sgRNAs designed to bind the CaMKK2 gene transcript. **G,** Representative blot showing CaMKK2 KO clones generated using sgRNAs #A, #B, #C, #D. **H,** Representative blot showing CaMKK2 KO clones generated using sgRNAs #1, #2, #3 (clones used for downstream experiments are marked with *). **I, J,** Representative images showing genetic ablation of CaMKK2 impaired wound healing ability of MDA-MB-231 cells *in vitro*. *P < 0.05, **P < 0.01, ***P < 0.005, *P* values were calculated using unpaired Student's *t* test.

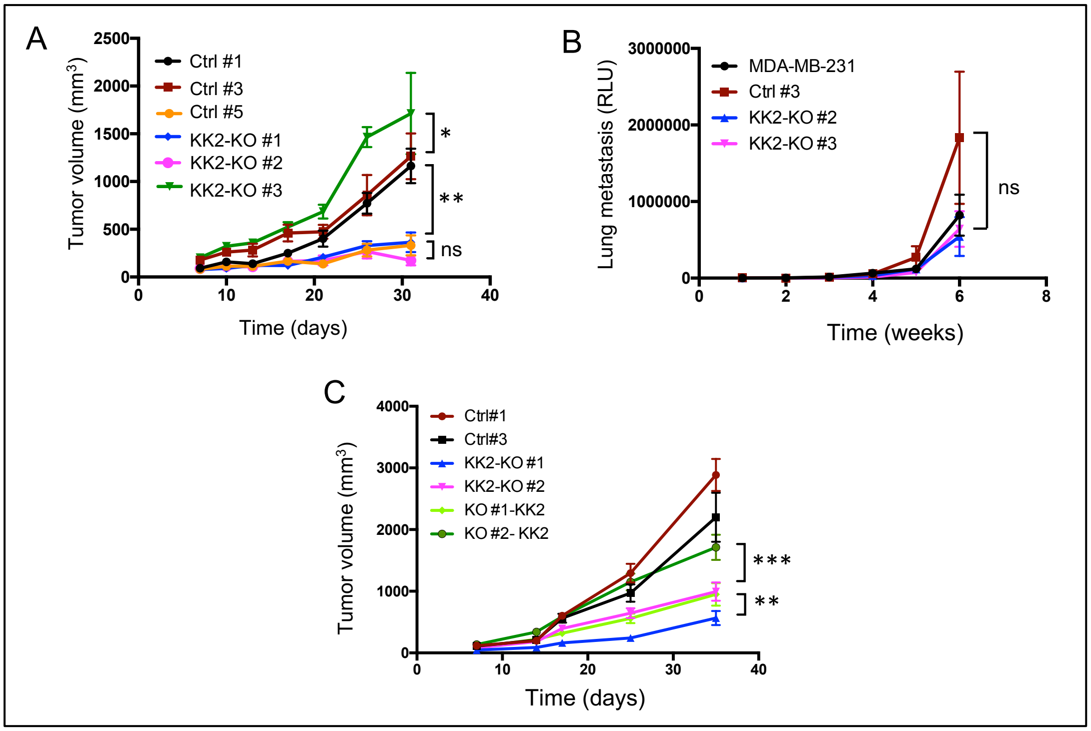


**Supplementary Figure 2. Genetic ablation of CaMKK2 does not affect primary tumor growth or the final colonization step of metastatic dissemination *in vivo*. Related to Figure 3. A,** Mice bearing the CaMKK2-KO tumors did not show consistent reduction in primary tumor growth rate as compared to mice bearing tumors from the control clones. 6-week old female nude mice were orthotopically injected in the mammary fat pad with three control clones (Ctrl #1, Ctrl #3, Ctrl #5) and three CaMKK2-KO (KK2-KO #1, KK2-KO #2, KK2-KO #3) clones. Tumor growth rate was monitored by measuring primary tumors with calipers twice weekly. **B,** Ablation of CaMKK2 did not impair the final seeding and colonization step of metastatic dissemination. Parental MDA-MB-231 cells, CRISPR control cells, and CaMKK2-KO cell clones (KK2-KO #1 and KK2-KO #2) were orthotopically injected into the tail vein of 6-week old female nude mice. Metastatic burden was monitored by BLI imaging. **C,** Addback of CaMKK2 expression in CaMKK2-KO clones #1 and #2 restored the reduced primary tumor growth rate. Two control cell clones (Ctrl #1, Ctrl #3), two CaMKK2-KO cell clones (KK2-KO #1, KK2-KO #2) and the CaMKK2 KO cell clones with re-expression of CaMKK2 (KO #1-KK2, KO #2-KK2) were orthotopically injected in the mammary fat pad of 6-week old female nude mice. Tumor growth rate was monitored by measuring primary tumors with calipers twice weekly. Data are expressed as the mean ± SEM. *$P < 0.05$, **$P < 0.01$, ***$P < 0.005$, and ****$P < 0.001$, by two-way ANOVA followed by Bonferroni's multiple-correction test.

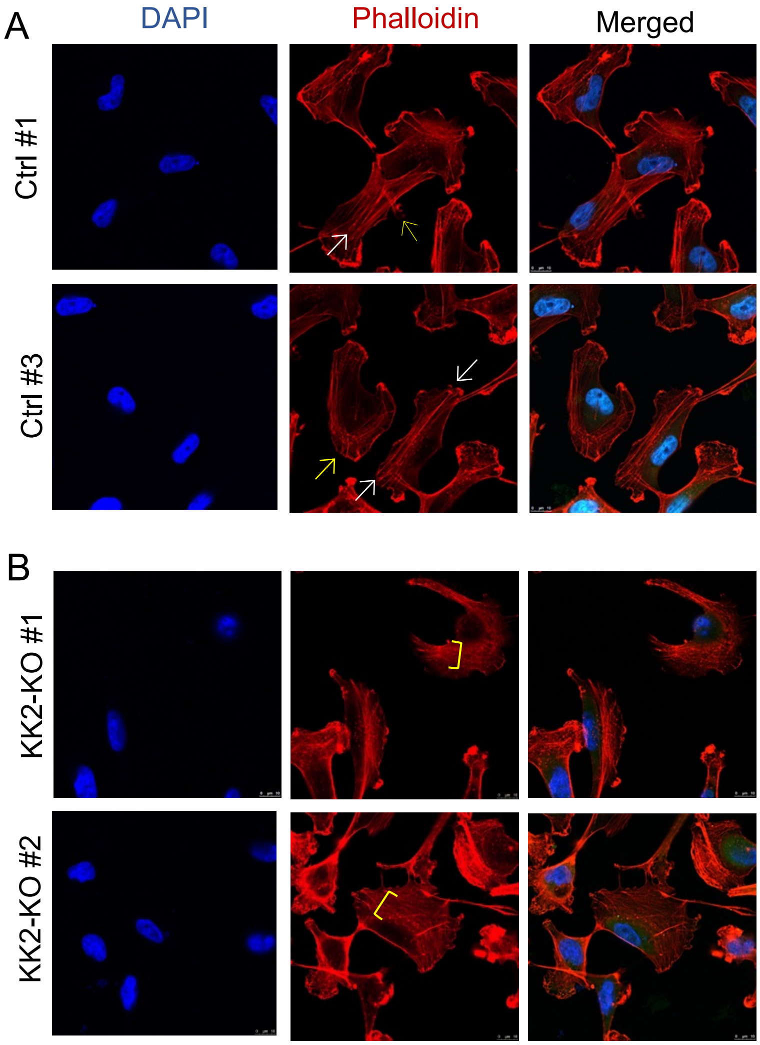


**Supplementary Figure 3. Representative images showing impaired cytoskeletal assembly in CaMKK2-KO cells as compared to control cells. Related to Figure 5.** Visualization of F-actin by phalloidin staining in CaMKK2-WT (Ctrl #1) and CaMKK2-KO (KK2-KO #1) cells revealed almost complete loss of thick ventral stress fibers in cells lacking CaMKK2 expression. Yellow arrows, dorsal stress fibers; white arrows, ventral stress fibers; yellow bracket, transverse arcs. The scale bar represents 10 µm.

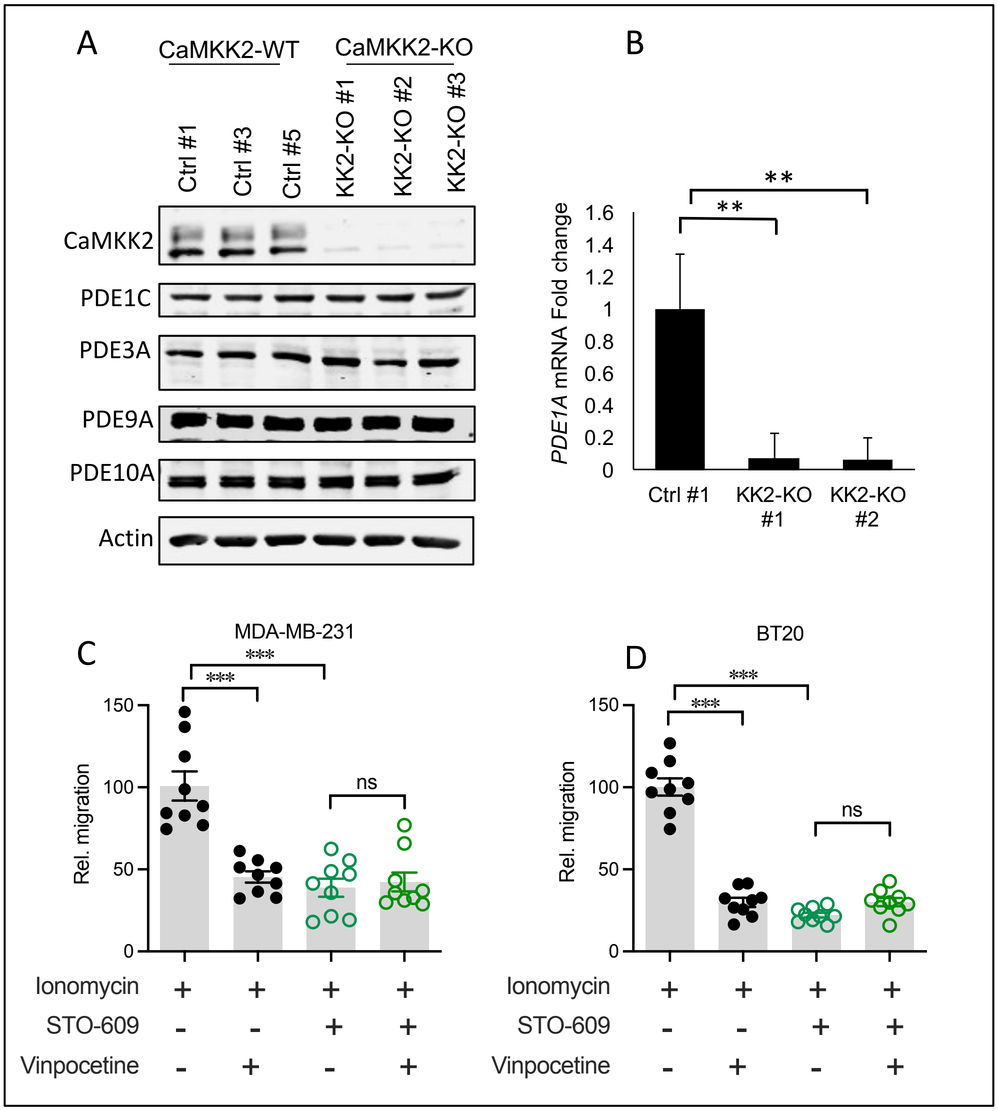


**Supplementary Figure 4. Pharmacological inhibition of PDE1 diminishes the migratory capacity of breast cancer cells. Related to Figure 7. A,** Representative blot showing that depletion of CaMKK2 does not impact the expression of cGMP specific PDEs other than PDE1A. Cell lysates were harvested and western immunoblots were used to analyze CaMKK2 and PDE expression levels. β-Actin was used as a loading control. This western immunoblot was performed once. **B**, Depletion of CaMKK2 results in reduced PDE1A expression at the transcriptional level, qRT-PCR profiling of the expression of *PDE1A* is shown. Data are plotted as mean ± SEM as representative results from two independent experiments. **C, D,** Inhibition of PDE1 with vinpocetine reduced the migratory capacity of parental MDA-MB-231 and BT-20 cells. Data are plotted as mean ± SEM; *n* = 4-5 random fields measurements from two individual transwells.

**Supplementary Video legends**

**Supplementary Video 1. Related to Figure 2K.** Control cells (Ctrl#1) were stained with Cell Brite Green and mixed with freshly isolated minced human breast tissue in a SWAT petridish (see Methods). Cancer cell movement was tracked by live cell imaging capturing fluorescence microscopy and bright field images every 5 minutes over 6 hours.

**Supplementary Video 2. Related to Figure 2K.** The movement of Control cells (Ctrl#3) stained with Cell Brite Green and mixed with minced human breast tissue in a SWAT petridish was tracked by capturing fluorescence microscopy and bright field images every 5 minutes over 6 hours.

**Supplementary Video 3. Related to Figure 2K.** CaMKK2-KO cells (KK2-KO#1) were stained with Cell Brite Green and mixed with freshly isolated minced human breast tissue in a SWAT petridish as described. Cancer cell movement over adipocytes was tracked by live cell imaging capturing fluorescence microscopy and bright field images every 5 minutes over 6 hours.

**Supplementary Video 4. Related to Figure 2K.** CaMKK2-KO cells (KK2-KO#2) stained with Cell Brite Green and mixed with freshly isolated minced human breast tissue in a SWAT petridish was visualized by live cell imaging. Cellular movement over adipocytes was tracked by capturing fluorescence microscopy and bright field images every 5 minutes over 6 hours.

**Supplementary Table 1**

Sequences of siRNAs

| | |
|---|---|
| siKK2 #1 | GGA CCA UCU GUA CAU GGU GUU CGA A |
| siKK2 #2 | GCU GAC UUU GGU GUG AGC AAU GAA U |

**Supplementary Table 2**

Sequences of *CAMKK2* sgRNAs

| | |
|---|---|
| sgRNA #A | CCC CAG CUC AUC CUG GGG GG |
| sgRNA #B | UAG CCA GCC CAG CAG CAA CC |
| sgRNA #C | GGG GCG GCC CGG UUG CUG CU |
| sgRNA #D | GCU AGA GAC ACA UGA UGA CA |
| sgRNA #1 | AGC UUG CGA CCG GAG AGG UG |
| sgRNA #2 | ACA CUC GGU GAG CAC AAU GA |
| sgRNA #3 | UGA AGG ACU CCA UGC CCA GG |
